# Sharing of Very Short IBD Segments between Humans, Neandertals, and Denisovans

**DOI:** 10.1101/003988

**Authors:** Gundula Povysil, Sepp Hochreiter

## Abstract

We analyze the sharing of very short identity by descent (IBD) segments between humans, Neandertals, and Denisovans to gain new insights into their demographic history. Short IBD segments convey information about events far back in time because the shorter IBD segments are, the older they are assumed to be. The identification of short IBD segments becomes possible through next generation sequencing (NGS), which offers high variant density and reports variants of all frequencies. However, only recently HapFABIA has been proposed as the first method for detecting very short IBD segments in NGS data. HapFABIA utilizes rare variants to identify IBD segments with a low false discovery rate.

We applied HapFABIA to the 1000 Genomes Project whole genome sequencing data to identify IBD segments that are shared within and between populations. Many IBD segments have to be old since they are shared with Neandertals or Denisovans, which explains their shorter lengths compared to segments that are not shared with these ancient genomes. The Denisova genome most prominently matches IBD segments that are shared by Asians. Many of these segments were found exclusively in Asians and they are longer than segments shared between other continental populations and the Denisova genome. Therefore, we could confirm an introgression from Deniosvans into ancestors of Asians after their migration out of Africa. While Neandertal-matching IBD segments are most often shared by Asians, Europeans share a considerably higher percentage of IBD segments with Neandertals compared to other populations, too. Again, many of these Neandertal-matching IBD segments are found exclusively in Asians, whereas Neandertal-matching IBD segments that are shared by Europeans are often found in other populations, too. Neandertal-matching IBD segments that are shared by Asians or Europeans are longer than those observed in Africans. These IBD segments hint at a gene flow from Neandertals into ancestors of Asians and Europeans after they left Africa. Interestingly, many Neandertal-and/or Denisova-matching IBD segments are predominantly observed in Africans — some of them even exclusively. IBD segments shared between Africans and Neandertals or Denisovans are strikingly short, therefore we assume that they are very old. Consequently, we conclude that DNA regions from ancestors of humans, Neandertals, and Denisovans have survived in Africans. As expected, IBD segments on chromosome X are on average longer than IBD segments on the autosomes. Neandertal-matching IBD segments on chromosome X confirm gene flow from Neandertals into ancestors of Asians and Europeans outside Africa that was already found on the autosomes. Interestingly, there is hardly any signal of Denisova introgression on the X chromosome.

## 1 Introduction

The recent advent of next generation sequencing technologies made whole genome sequencing of thousands of individuals feasible (52). These sequencing techniques have been extended to allow sequencing of ancient DNA which enabled researchers to assemble the DNA of hominid individuals that lived ten thousands of years ago (16, 35, 39, 41). These advances in biotechnology could help to answer one of the most fundamental questions of humanity: “Where do we come from?”

In 2012, the 1000 Genomes Project Consortium (53) published first results on the genomes of more than thousand individuals from 14 populations stemming from Europe, East Asia, Africa, and the Americas. Reviewing the differences in genetic variation within and between populations, the authors found evidence of past events such as bottlenecks or admixtures, but also regions which seemed to be under a strong selective pressure. Gravel et al. (15) used the sequence data of the individuals from the Americas of the 1000 Genomes Project to quantify the contributions of European, African, and Native American ancestry to these populations, to estimate migration rates and timings, as well as to develop a demographic model.

Green et al. (16) were the first to analyze a draft sequence of the Neandertal genome derived from the bones of three individuals from Vindija Cave in Croatia that died about 40,000 years ago. They found that non-African individuals share more alleles with the Neandertal genome than sub-Saharan Africans. Until then evidence from mtDNA (9, 28, 47) and the Y chromosome (27) suggested that Neandertals lived isolated in Europe and Asia until they were replaced by anatomically modern humans. Although this theory still holds if differences in allele sharing are attributed to the existence of an ancient population substructure within Africa (10, 11, 33), ancient admixture events between Neandertals and anatomically modern humans outside of Africa are considered as a more plausible explanation (16, 32, 44, 61). Further studies (35, 60) reported more Neandertal DNA preserved in modern East Asians than Europeans, hinting at some admixture after the separation of ancestors of Europeans and Asians. Prüfer et al. (39) confirmed higher rates of sharing for non-Africans using a high-quality genome sequence of a Neandertal from Denisova Cave in the Altai Mountains of Siberia. The authors further report that populations in Asia and America have more regions of Neandertal origin than populations in Europe.

A bone of a sister group of Neandertals, named Denisovans after the cave in the Altai Mountains of Siberia, was sequenced first at low (41) and later at high coverage (35). Studies on the low coverage draft sequence, as well as on the high coverage sequence, reported contributions from Denisovans to the gene pool of present-day individuals in Southeast Asia and Oceania (35, 41, 42, 48). Additionally to the clear signal of the Denisova genome found in Oceanians, Prüfer et al. (39) detected some regions of Denisovan origin in East and Southeast Asians and also in populations of the Americas, but only very few in Europeans. The authors further report gene flow from Neandertals and an unknown archaic population into Denisovans indicating that interbreeding among distinct hominid populations was more common than previously thought.

Using the high coverage Altai Neandertal genome Sankararaman et al. (43) screened the 1000 Genomes Project data for Neandertal ancestry with the genomes of the West African Yoruba individuals from Ibadan, Nigeria, as a reference panel that are assumed to harbor no Neandertal ancestry. They again found more Neandertal ancestry in East-Asians than in Europeans, but also very low levels in African Luhya individuals from Webuye, Kenya. The authors further looked for regions in the genome with very high or very low Neandertal ancestry that both might be due to selective pressure. Vernot and Akey (58) searched for signatures of introgression in the sequences of Asian and European individuals of the 1000 Genomes Project and compared them with the Altai Neandertal genome. They confirmed more Neandertal DNA in East Asians than Europeans and looked for different levels of introgression along the genome.

In this study we propose to use short segments of identity by descent (IBD) to infer the population structure of humans and to gain insights into the genetic relationship of humans, Neandertals and Denisovans.

### 1.1 IBD for Inferring Population Structure

A DNA segment is *identical by state (IBS)* in two or more individuals if they all have identical nucleotide sequences in this segment. An IBS segment is *identical by descent (IBD)* in two or more individuals if they have inherited it from a common ancestor, that is, the segment has the same ancestral origin in these individuals. Rare variants can be utilized for distinguishing IBD from IBS without IBD because independent origins are highly unlikely for such variants. In other words, IBS generally implies IBD for rare variants, which is not true for common variants (49, Ch. 15.3, p. 441).

IBD detection methods have already been successfully used for inferring population structure. Gusev et al. (17) looked for long IBD segments shared within and between populations to estimate the demographic history of Ashkenazi Jewish individuals. Using similar models Palamara et al. (38) and Carmi et al. (7) reconstructed the demographic history of Ashkenazi Jewish and Kenyan Maasai individuals. Botigué et al. (3) confirmed gene flow from North Africans into Southern Europeans via patterns of long shared IBD segments. Ralph and Coop (40) tried to quantify the recent shared ancestry of different European populations by looking for long segments of shared DNA. Gravel et al. (15) similarly tried to draw conclusions of the genetic history of populations in the Americas using the respective data of the 1000 Genomes Project.

Except for Gravel et al. all of these studies were performed on SNP microarray data as IBD segments could not reliably be detected in large sequencing data. Sequencing data has higher marker density and also captures rare variants in contrast to SNP microarray data, therefore it would allow for a finer resolution of the length of IBD segments. Furthermore, all previous studies based on microarrays were limited to long IBD segments that stem from a very recent common ancestor. However, shorter IBD segments would convey information about events farther back in time because the shorter IBD segments are, the older they are assumed to be. Therefore, existing studies were not able to resolve demographic histories at a fine scale and very far back into the past.

### 1.2 HapFABIA for Extracting Short IBD Segments

We recently developed HapFABIA (21) (see http://dx.doi.org/10.1093/nar/gkt1013) to identify very short segments of identity by descent (IBD) that are tagged by rare variants (the so called tagSNVs) in large sequencing data. HapFABIA identifies **100 times smaller IBD segments than current state-of-the-art methods: 10 kbp for HapFABIA vs. 1 Mbp for state-of-the-art methods**. HapFABIA utilizes rare variants (*≤*5% MAF) to distinguish IBD from IBS without IBD because independent origins of rare minor alleles are highly unlikely (49, Ch. 15.3,p. 441). More importantly, rare variants make juxtapositions of smaller IBD segments unlikely which prevents the summary of several small IBD segment into one long IBD segment. Consequently, the length of IBD segments is estimated more accurately than with previous methods.

In experiments with artificial, simulated, and real genotyping data HapFABIA outperformed its competitors in detecting short IBD segments (21). HapFABIA is based on biclustering (22) which in turn uses machine learning techniques derived from maximizing the posterior in a Bayes framework (8, 23, 24, 25, 34, 46, 50, 51).

HapFABIA is designed to detect short IBD segments in genotype data that were obtained from next generation sequencing (NGS), but can also be applied to DNA microarray data. Especially in NGS data, HapFABIA exploits rare variants for IBD detection. Rare variants convey more information on IBD than common variants, because random minor allele sharing is less likely for rare variants than for common variants (5). In order to detect short IBD segments, both the information supplied by rare variants and the information from IBD segments that are shared by more than two individuals should be utilized (5). HapFABIA uses both. The probability of randomly sharing a segment depends

- (a) on the allele frequencies within the segment, where lower frequency means lower probability of random sharing, and
- (b) on the number of individuals that share the allele, where more individuals result in lower probability of random segment sharing.

The shorter the IBD segments, the higher the likelihood that they are shared by more individuals. A segment that contains rare variants and is shared by more individuals has higher probability of representing IBD (19, 31). These two characteristics are our basis for detecting short IBD segments by HapFABIA.

## 2 IBD Segments in the 1000 Genomes Data

### 2.1 The 1000 Genomes Data

We used HapFABIA to extract short IBD segments from the 1000 Genomes Project genotyping data (53), more specifically, the phase 1 integrated variant call set (version 3) containing phased genotype calls for SNVs, short indels, and large deletions. This data set consists of 38.2M SNVs, 1.4M short indels, and 14k large deletions from 1,092 individuals (246 Africans, 181 Admixed Americans, 286 East Asians, and 379 Europeans). We removed 36 individuals because they showed cryptic relatedness to others. The final data set consisted of 1,056 individuals (229 Africans, 175 Admixed Americans, 276 East Asians, and 376 Europeans).

Chromosomes 1-22 contain 37,802,491 SNVs and short indels that are on average 73 bp apart and have an average minor allele frequency (MAF) of 0.05. 30,541,037 (80.8%) variants are rare (MAF *≤* 0.05 including privates), 8,289,575 (21.9%) are private (minor allele is observed only once), and 7,261,454 (19.2%) are common (MAF *>* 0.05).

Chromosome X contains 1,154,686 SNVs and short indels that are on average 104 bp apart and have an average MAF of 0.06. 900,453 (78.0%) variants are rare (MAF *≤* 0.05 including privates), 235,179 (20.4%) are private (minor allele is observed only once), 254,233 (22.0%) are common (MAF *>* 0.05).

All chromosomes were divided into intervals of 10,000 SNVs with adjacent intervals overlapping by 5,000 SNVs. After removing common and private SNVs, we applied HapFABIA with default parameters to these intervals ignoring phase information since previous analyses revealed many phasing errors.

In the following, we separately report findings for autosomes and chromosome X, since inheritance, recombination and mutation rates, influences of natural selection, etc. differ between chromosome X and the autosomes (45). Chromosome Y is not considered as it is too small to yield reliable results concerning IBD segments, hardly has recombinations, and differs from other chromosomes as it is inherited only by males from males.

### 2.2 Summary Statistics of the Detected Short IBD Segments

HapFABIA found 1,348,320 different very short IBD segments on the autosomes. These contained 8,081,564 rare tagSNVs, which amounts to 26.5% of the rare variants (36.3% if private SNVs are excluded) and 21.4% of all SNVs. The distance between centers of IBD segments had a median of 1 kbp and a mean of 2.1 kbp and ranged from 0 (overlapping IBD segments) to several Mbp. The number of individuals that shared the same IBD segment was between 2 and 138, with a median of 5 and a mean of 11. IBD segments were tagged by 8 to 271 tagSNVs, with a median of 13 and a mean of 19. The length of IBD segments ranged from 15 base pairs to 21 Mbp, with a median of 24 kbp and a mean of 26 kbp. IBD lengths were computed as described in Section 4.1, to match the assumptions for the distribution of IBD segment lengths as derived in other publications (4, 17, 55, 56).

On chromosome X, HapFABIA detected 43,069 different IBD segments containing 305,028 rare tagSNVs, which amounts to 33.9% of the rare variants (45.8% if private SNVs are excluded) and 26.4% of all SNVs. The distance between centers of IBD segments had a median of 1.8 kbp and a mean of 3.6 kbp and ranged from 0 (overlapping IBD segments) to several Mbp. The number of individuals that shared the same IBD segment was between 2 and 96, with a median of 6 and a mean of 11. IBD segments were tagged by 8 to 255 tagSNVs, with a median of 15 and a mean of 21. The length of IBD segments ranged from 44 base pairs to 3.2 Mbp, with a median of 33 kbp and a mean of 36 kbp. On average, IBD segments on chromosome X were longer than IBD segments found on the autosomes.

## 3 Sharing of IBD Segments Between Populations and With Ancient Genomes

### 3.1 Sharing of IBD Segments Between Human Populations

We were interested in the distribution of IBD segments among different populations. The main continental population groups are Africans (AFR), Asians (ASN), Europeans (EUR), and Admixed Americans (AMR), where AMR consist of Colombian, Puerto Rican, and Mexican individuals. Table 1 lists the number and percentage of IBD segments on the autosomes that are shared between particular continental populations. The vast majority (1,294,240) of the detected IBD segments (1,348,320) is shared by Africans (at least one African possesses the segment), of which 837,769 are exclusively found in Africans. Only 118,775 and 54,180 IBD segments are shared by Europeans and Asians, respectively. 7,621 IBD segments are exclusively found in Europeans and 15,438 exclusively in Asians. Admixed Americans share 2,744 IBD segments with Asians, but 13,793 with Europeans. Gravel et al. (15) reported recently that Admixed Americans, especially Colombians and Puerto Ricans, show a large proportion of European ancestry, which is also reflected in our results. If we additionally consider sharing with AFR, we obtain similar figures that are again consistent with the AMR admixture: 58,114 IBD segments have AFR/AMR/EUR sharing while only 6,715 IBD segments have AFR/AMR/ASN sharing. 14,179 IBD segments are shared by individuals from all continental populations. According to results of the 1000 Genomes Project Consortium (53), individuals with African ancestry carry much more rare variants than those of European or Asian ancestry supporting our finding that most IBD segments are shared by Africans. We found that few IBD segments are shared between two continental populations (Table 1 “Pairs of Populations”) confirming recently published results (14, 53). The relatively large number of shared IBD segments between Africans and Europeans was due to many shared IBD segments between the AFR sub-group ASW (Americans with African ancestry from SW US) and Europeans. This tendency was also observed in the 1000 Genomes Project via the fixation index *F*_ST_ estimated by Hudson ratio of averages and via shared haplotype length around *f*_2_ variants (53). The high content of European DNA segments in ASW is consistent with the finding that in African Americans a median proportion of 18.5% is European (6).

**Table 1:**
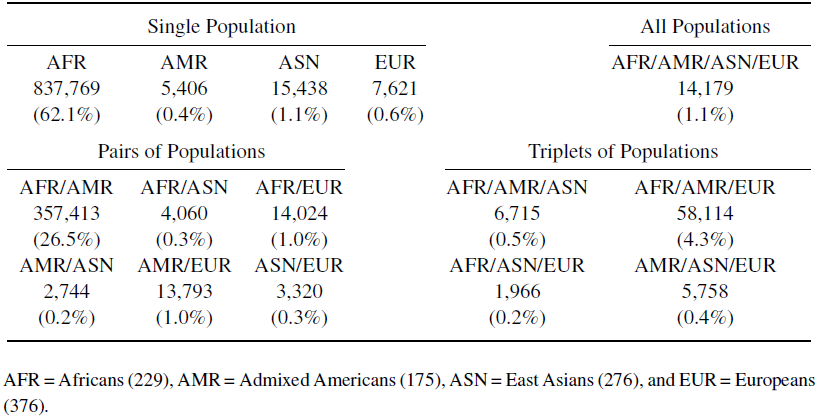
Number and percentage of IBD segments on the autosomes that are shared by particular continental populations.

| Single Population |  |  |  | All Populations |
| --- | --- | --- | --- | --- |
| AFR | AMR | ASN | EUR | AFR/AMR/ASN/EUR |
| 837,769 | 5,406 | 15,438 | 7,621 | 14,179 |
| (62.1%) | (0.4%) | (1.1%) | (0.6%) | (1.1%) |
| Pairs of Populations |  |  | Triplets of Populations |  |
| AFR/AMR | AFR/ASN | AFR/EUR | AFR/AMR/ASN | AFR/AMR/EUR |
| 357,413 | 4,060 | 14,024 | 6,715 | 58,114 |
| (26.5%) | (0.3%) | (1.0%) | (0.5%) | (4.3%) |
| AMR/ASN | AMR/EUR | ASN/EUR | AFR/ASN/EUR | AMR/ASN/EUR |
| 2,744 | 13,793 | 3,320 | 1,966 | 5,758 |
| (0.2%) | (1.0%) | (0.3%) | (0.2%) | (0.4%) |

AFR = Africans (229), AMR = Admixed Americans (175), ASN = East Asians (276), and EUR = Europeans (376).

Table 2 lists the number and percentage of IBD segments on chromosome X that are shared between particular continental populations. Again, the vast majority (41,636) of the detected IBD segments (43,069) is shared by Africans (at least one African possesses the segment), of which 23,087 are exclusively found in Africans. Only 4,226 and 1,958 IBD segments are shared by Europeans and Asians, respectively. 251 IBD segments are exclusively found in Europeans and 335 exclusively in Asians. Admixed Americans share 96 IBD segments with Asians, but 339 with Europeans. 2,045 IBD segments have AFR/AMR/EUR sharing while only 305 IBD segments have AFR/AMR/ASN sharing. 696 IBD segments are shared by individuals from all continental populations.

**Table 2:** Number and percentage of IBD segments on chromosome X that are shared by particular continental populations.

| Single Population |  |  |  | All Populations |
| --- | --- | --- | --- | --- |
| AFR | AMR | ASN | EUR | AFR/AMR/ASN/EUR |
| 23,087 | 183 | 335 | 251 | 696 |
| (53.6%) | (0.4%) | (0.8%) | (0.5%) | (1.6%) |
| Pairs of Populations |  |  | Triplets of Populations |  |
| AFR/AMR | AFR/ASN | AFR/EUR | AFR/AMR/ASN | AFR/AMR/EUR |
| 14,633 | 204 | 573 | 305 | 2,045 |
| (34.0%) | (0.5%) | (1.3%) | (0.7%) | (4.8%) |
| AMR/ASN | AMR/EUR | ASN/EUR | AFR/ASN/EUR | AMR/ASN/EUR |
| 96 | 339 | 86 | 93 | 143 |
| (0.2%) | (0.8%) | (0.2%) | (0.2%) | (0.3%) |

AFR = Africans (229), AMR = Admixed Americans (175), ASN = East Asians (276), and EUR = Europeans (376).

Overall, we conclude that IBD segments that are shared across continental populations, in particular by Africans, date back to a time before humans moved out of Africa. Consequently, the rare variants that tag these short IBD segments arose before this time. See Section A, for a discussion of the question whether rare variants are recent or old.

### 3.2 Sharing of IBD Segments Between Human and Ancient Genomes

Since short IBD segments are thought to be very old, we wondered whether some IBD segments match bases of ancient genomes, such as Neandertal and Denisova. Ancient short IBD segments may reveal gene flow between ancient genomes and ancestors of modern humans and, thereby, may shed light on different out-of-Africa hypotheses (2). 31X coverage whole genome sequencing data for the Denisova, as well as, the Altai Neandertal genome of 52X coverage were provided by the Max Planck Institute for Evolutionary Anthropology (35, 39). Considering only the variants of the 1000 Genomes Project, 0.5% of the Denisova bases and 0.3% of the Neandertal bases were not determined, 92.0% and 91.3%, respectively, matched bases of the human reference, and 8% and 9%, respectively, matched either the human minor allele or were different from human alleles. As additional information in the 1000 Genomes Project data, bases of the reconstructed common ancestor of human, chimpanzee, gorilla, orang utan, macaque, and marmoset genomes were given.

#### 3.2.1 IBD Sharing Between Human Populations and Ancient Genomes

We tested whether IBD segments that match particular ancient genomes to a large extent are found more often in certain populations than expected randomly. For each IBD segment, we computed two values: The first value was the proportion of tagSNVs that match a particular ancient genome, which we call “genome proportion” of an IBD segment (e.g. “Denisova proportion”). The second value was the proportion of individuals that possess an IBD segment and are from a certain population as opposed to the overall number of individuals that possess this IBD segment. We call this value the “population proportion” of an IBD segment (e.g. “Asian proportion”). Consider the following illustrative examples. If an IBD segment has 20 tagSNVs of which 10 match Denisova bases with their minor allele, then we obtain 10/20 = 0.5 = 50% as the Denisova proportion. If an IBD segment is observed in 6 individuals of which 4 are Africans and 2 Europeans, then the African proportion is 4/6 = 0.67 = 67% and the European proportion is 2/6 = 0.33 = 33%. A correlation between a genome proportion and a population proportion would indicate that this ancient genome is overrepresented in this specific population. Often human IBD segments that are found in ancient genomes have already been present in the ancestral genome of humans and other primates. These IBD segments would confound the interbreeding analysis based on IBD sharing between modern human populations and Neandertals or Denisovans. Therefore, we removed those IBD segments from the data before the following analysis.

For the autosomes, Pearson’s product moment correlation test and Spearman’s rank correlation test both showed highly significant correlations between the Denisova genome and Asians, the Denisova genome and Europeans, the Neandertal genome and Asians, and the Neandertal genome and Europeans.

Figure 1 shows Pearson’s correlation coefficients for the correlation between population proportion and the Denisova genome proportion. Asians have a significantly larger correlation with the Denisova genome than other populations (*p*-value *<* 5e-324). Many IBD segments that match the Denisova genome are exclusively found in Asians, which has a large effect on the correlation coefficient. Europeans have still a significantly larger correlation with the Denisova genome than the average (*p*-value *<* 5e-324). Surprisingly, Mexicans (MXL) have a high correlation with the Denisova genome too, while Iberians (IBS) have a low correlation compared to other Europeans. These correlations may be explained through the fact that of all European populations Iberians show the highest rates of African gene flow (1, 3, 36) whereas Mexicans show a high proportion of Native American ancestry which in turn might also reflect gene flow from Asia (15, 37).

**Figure 1:**
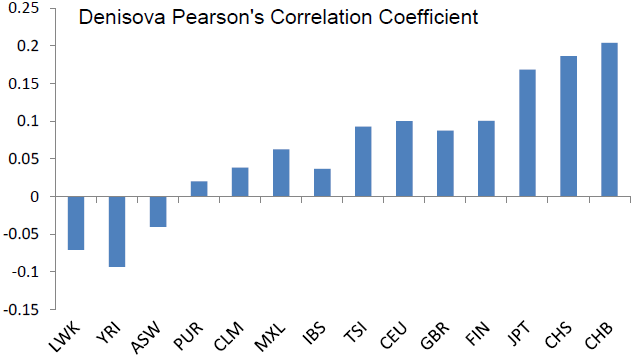
Pearson’s correlation between population proportions and the Denisova genome proportion. Asians have a significantly larger correlation with the Denisova genome than other populations. Many IBD segments that match the Denisova genome are exclusively found in Asians. Europeans still have a significantly larger correlation with the Denisova genome than the average. Mexicans (MXL) have also a surprisingly high correlation with the Denisova genome while Iberians (IBS) have a low correlation compared to other Europeans.

Figure 2 shows Pearson’s correlation coefficients for the correlation between population proportion and the Neandertal genome proportion. Europeans and Asians have a significantly larger correlation with the Neandertal genome than other populations (*p*-value *<* 5e-324). Asians have slightly higher correlation coefficients than Europeans. Again, Mexicans (MXL) have surprisingly high correlation with the Neandertal genome while Iberians (IBS) have a low correlation compared to other Europeans.

**Figure 2:**
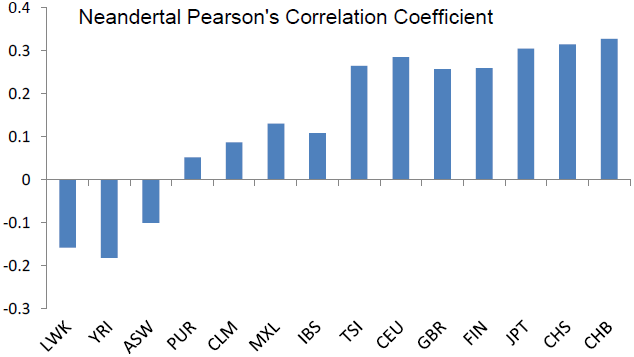
Pearson’s correlation between population proportions and the Neandertal genome proportion. Europeans and Asians have significantly larger correlations with the Neandertal genome than other populations. Asians have slightly higher correlation coefficients than Europeans. Again, Mexicans (MXL) have a surprisingly high correlation with the Neandertal genome while Iberians (IBS) have a low correlation compared to other Europeans.

Moving on to chromosome X, the results for the Neandertal genome are similar to the autosomes while findings concerning the Denisova genome are different. For the Neandertal genome, significant correlations could be found with Asians and Europeans, however they are smaller than on the autosomes. These correlations confirm the results from Sankararaman et al. (43) who also found lower Neandertal ancestry on chromosome X. Figure 3 shows Pearson’s correlation coefficients for the correlation between population proportions and the Neandertal genome proportion on chromosome X. Europeans and Asians have a significantly larger correlation with the Neandertal genome than other populations (*p*-value *<* 5e-324). Asians have slightly higher correlation coefficients than Europeans. Again, Iberians (IBS) have a very low correlation compared to other Europeans.

**Figure 3:**
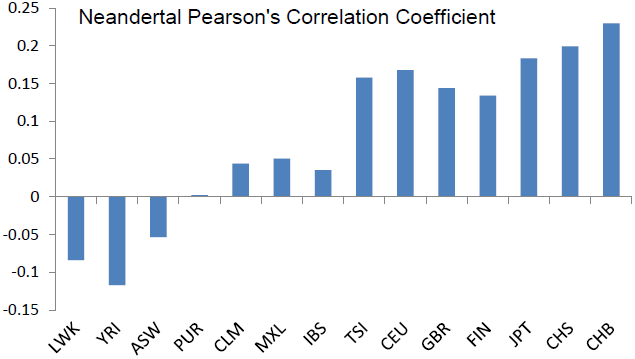
Pearson’s correlation between population proportions and the Neandertal genome proportion on chromosome X. Europeans and Asians have significantly larger correlations with the Neandertal genome than other populations. Asians have slightly higher correlation coefficients than Europeans. Again, Iberians (IBS) have a very low correlation compared to other Europeans. All correlations are weaker than on the autosomes.

Surprisingly, for the Denisova genome correlations to each of the tested present day populations were similarly low for the X chromosome. The small correlation coefficients indicate that almost no Denisova DNA segments can be found on the X chromosome compared to autosomes.

These findings can be explained either by the fact that Denisova-DNA never entered the human X chromosome outside Africa or on this chromosome the Denisova sequence got lost in modern humans. The former explanation would indicate that male Denisovans primarily fathered male offspring with human females. The latter explanation would hint at some selective pressure which is also assumed to be the reason for lower Neandertal ancestry on chromosome X (43). Meyer et al. (35) reported similar explanations for their finding that Papuans share more alleles with the Denisova on the autosomes than on chromosome X.

However, correlation tests are sensitive to accumulations of minor effects. Therefore, we focused subsequently on strong effects, i.e. large values of genome proportions and large values of population proportions.

We define an IBD segment to match a particular ancient genome if the genome proportion is 30% or higher. Only 8% of the Denisova and 9% of the Neandertal bases (about 10% of the called bases) match the minor allele of the human genome on average. Therefore, we require an odds ratio of 3 to call an IBD segment to match an ancient genome. We found many more IBD segments that match the Neandertal or the Denisova genome than expected randomly. This again supports the statement that the detected short IBD segments are old and some of them date back to times of the ancestors of humans, Neandertals, and Denisovans. IBD segments that match the Denisova genome often match the Neandertal genome, too, thus these segments cannot be attributed to either one of these genomes. Therefore, we introduce the “Archaic genome” (genome of archaic hominids ancestral to Denisova and Neandertal), to which IBD segments are attributed if they match both the Denisova and the Neandertal genome.

To test for an enrichment of Denisova-, Neandertal-, or Archaic-matching IBD segments in different subpopulations, we used Fisher’s exact test for count data. Again, we first excluded those IBD segments that were also present in the reconstructed ancestral genome of humans and other primates. IBD segments were then classified as being shared by at least one individual of a certain population or not and either matching or not matching the ancient genome.

Figure 4 shows the odds scores of the Fisher’s exact tests for an enrichment of Denisova-matching IBD segments in different populations. As expected, Asians show the highest odds for IBD segments matching the Denisova genome (odds ratio of 5.83 and *p*-value *<* 5e-324), while Africans have the lowest odds (odds ratio of 0.24 and *p*-value *<* 5e-324). Surprisingly, Admixed Americans show more sharing than Europeans (odds ratio of 2.61 vs. 2.26), although the difference is not very prominent. In general these results reflect previous findings. Meyer et al. (35) and Prüfer et al. (39) noted that Denisovans show more sharing with modern East Asians and South Americans (called Admixed Americans here) than with Europeans. Within the Asian populations Han Chinese from Beijing (CHB) have slightly higher odds for matching the Denisova genome than Han Chinese from South (CHS), while Japanese (JPN) have the lowest odds (Figure 5). Meyer et al. (35) also found higher levels of sharing for individuals from North China compared to those of the south, while Skoglund et al. (48) report a particularly high affinity between Southeast Asians and the Denisova genome. Within the European populations we see the highest odds for Finnish individuals (FIN) followed by Toscani in Italy (TSI), Utah residents with ancestry from northern and western Europe (CEU), and British from England and Scotland (GBR). Iberians from Spain (IBS) show the lowest levels of sharing, but all odds are between 1.87 and 2.56 (Figure 6).

**Figure 4:**
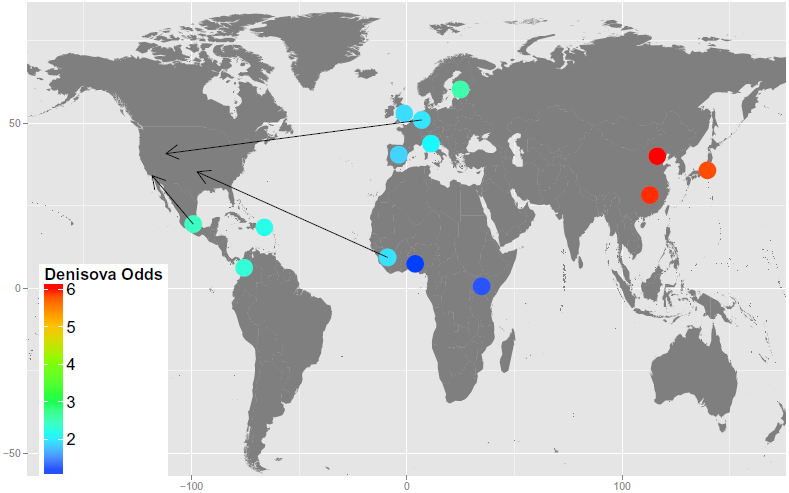
Odds scores of Fisher’s exact test for an enrichment of Denisova-matching IBD segments in different populations are represented by colored dots. The arrows point from the region the populations stem from to the region of sample collection. IBD segments that are shared by Asians match the Denisova genome significantly more often than IBD segments that are shared by other populations (red dots). Africans show the lowest matching with the Denisova genome (dark blue dots). Surprisingly, Admixed Americans have slightly higher odds for Denisova sharing than Europeans (turquoise vs. light blue dots).

**Figure 5:**
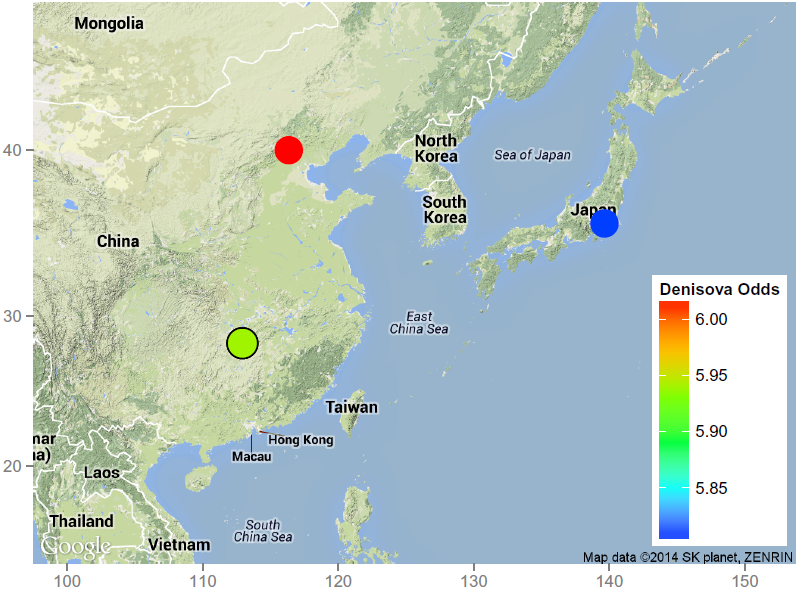
Odds scores of Fisher’s exact test for an enrichment of Denisova-matching IBD segments in Asian populations are represented by colored dots. Within the Asian populations Han Chinese from Beijing (CHB) have slightly higher odds for matching the Denisova genome than Han Chinese from South (CHS) (red dot vs. green dot), while Japanese (JPN) have the lowest odds (blue dot).

**Figure 6:**
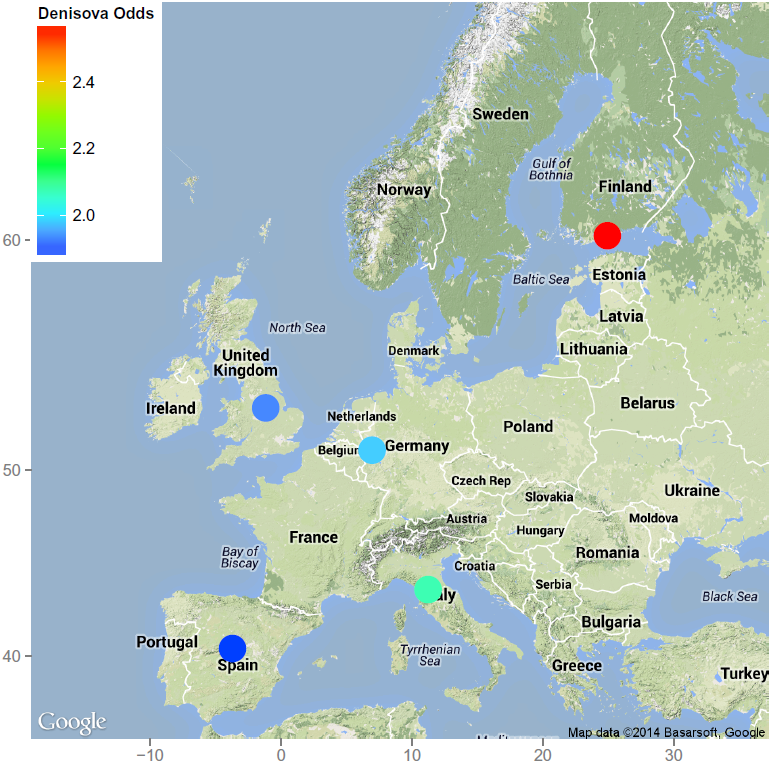
Odds scores of Fisher’s exact test for an enrichment of Denisova-matching IBD segments in European populations are represented by colored dots. Utah residents with ancestry from northern and western Europe (CEU) are symbolized by a dot in Central Europe. Within the European populations we see the highest odds (red dot) for Finnish individuals (FIN) followed by Toscani in Italy (TSI), Utah residents with ancestry from northern and western Europe (CEU), and British from England and Scotland (GBR). Iberians from Spain (IBS) show the lowest levels of sharing (dark blue dot), but all odds are between 1.87 and 2.56.

Figure 7 shows the odds scores of the Fisher’s exact tests for an enrichment of Neandertal-matching IBD segments in different populations. As expected, Asians have the highest odds for IBD segments matching the Neandertal genome (odds ratio of 22.16 and *p*-value *<* 5e-324), while Africans have again the lowest odds (odds ratio of 0.04 and *p*-value *<* 5e-324). In contrast to the Denisova results, here Europeans show clearly more matching with the Neandertal genome than Admixed Americans (odds ratio of 12.98 vs. 3.32). Other studies (16, 32) have also reported significantly more sharing between Neandertals and non-Africans than with Africans. Wall et al. (60) noted that Neandertals contributed more DNA to modern East Asians than to Europeans. Within the Asian populations Han Chinese from Beijing (CHB) and Japanese (JPN) have slightly higher odds for matching the Neandertal genome than Han Chinese from South (CHS). (Figure 8). Within the European populations a north to south decline can be seen with Finnish (FIN) having the highest odds for matching the Neandertal genome followed by British from England and Scotland (GBR) and Utah residents with ancestry from northern and western Europe (CEU). Toscani in Italy (TSI) and Iberians from Spain (IBS) show the lowest levels of matching the Neandertal genome (Figure 9). One explanation for the latter two low odds for Neandertal matching is that according to Ralph and Coop (40) Iberians and Italians show reduced rates of shared ancestry compared to the rest of Europe. Also Botigué et al. (3) found higher IBD sharing between North Africans and individuals from Southern Europe which would decrease the amount of DNA sharing with Neandertals. Sankararaman et al. (43) achieved a similar ranking for Neandertal ancestry using a different approach, but stated that because of the high standard deviation among individuals from the same population, small differences may be due to statistical noise. The same holds true for our results since a single individual with very high Neandertal ancestry can lead to inflated odds for a population. Nevertheless, since all Asian populations show especially high odds, we can exclude that all results are just due to noise.

**Figure 7:**
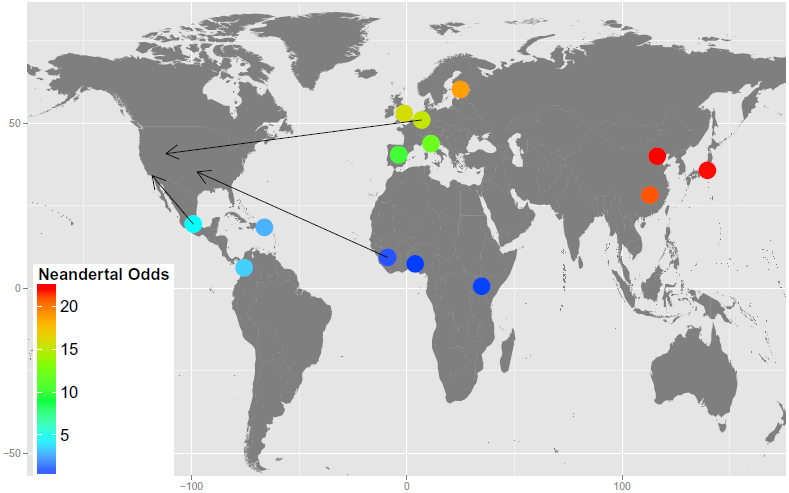
Odds scores of Fisher’s exact test for an enrichment of Neandertal-matching IBD segments in different populations are represented by colored dots. The arrows point from the region the populations stem from to the region of sample collection. IBD segments that are shared by Asians match the Neandertal genome significantly more often than IBD segments that are shared by other populations (red dots). Africans show the lowest matching with the Neandertal genome (dark blue dots). Europeans show more matching than Admixed Americans (green and orange vs. light blue dots).

**Figure 8:**
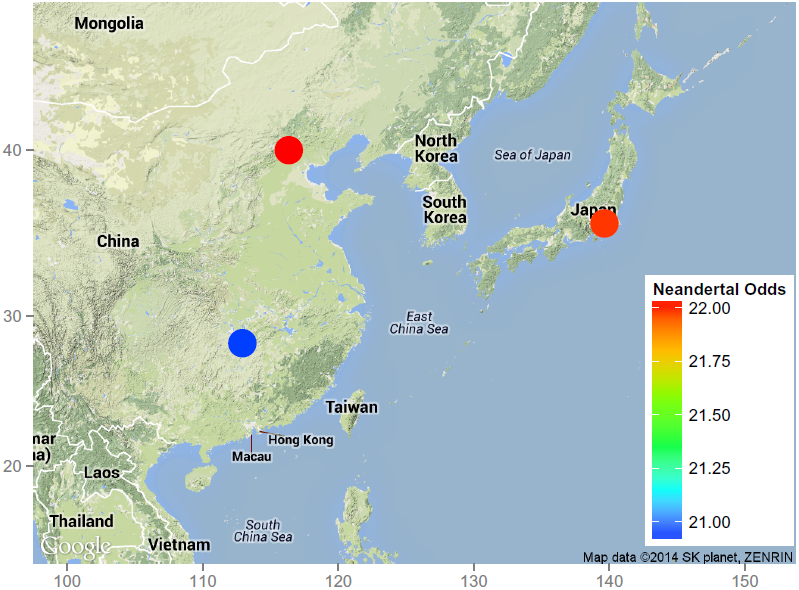
Odds scores of the Fisher’s exact tests for an enrichment of Neandertal-matching IBD segments in Asian populations are represented by colored dots. Within the Asian populations Han Chinese from Beijing (CHB) and Japanese (JPN) have slightly higher odds for matching the Neandertal genome than Han Chinese from South (CHS) (red dots vs. blue dot).

**Figure 9:**
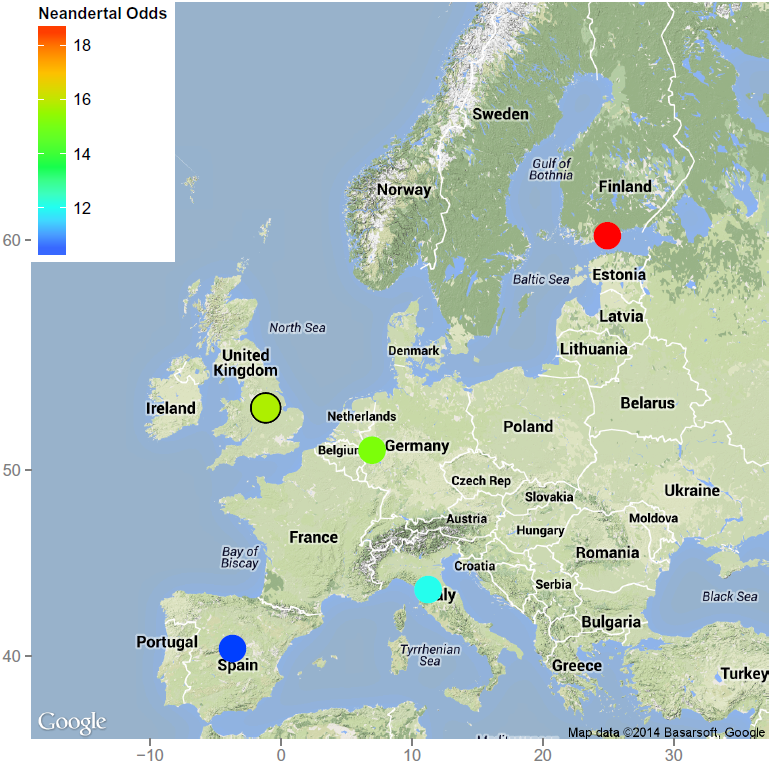
Odds scores of the Fisher’s exact tests for an enrichment of Neandertal-matching IBD segments in European populations are represented by colored dots. Utah residents with ancestry from northern and western Europe (CEU) are symbolized by a dot in Central Europe. Within the European populations a north to south decline can be seen with Finnish (FIN) having the highest odds for matching the Neandertal genome (red dot) followed by British from England and Scotland (GBR) and Utah residents with ancestry from northern and western Europe (CEU) (green dots). Toscani in Italy (TSI) and Iberians from Spain (IBS) show the lowest levels of matching (blue dots).

Figure 10 shows the odds scores of the Fisher’s exact tests for an enrichment of Archaic-matching IBD segments in different populations. As expected, Asians again show the highest odds for IBD segments matching both the Denisova and the Neandertal genome (odds ratio of 13.78 and *p*-value *<* 5e-324), while African populations have the lowest odds (odds ratio of 0.11 and *p*-value *<* 5e-324). Similar to the Neandertal results, Europeans show clearly more matching with the Archaic genome than Admixed Americans (odds ratio of 7.70 vs. 3.63). Within the Asian populations Japanese (JPN) have slightly higher odds for matching the Archaic genome than Han Chinese from South (CHS), while Han Chinese from Beijing (CHB) lie in between, but the differences are very small (Figure 11). Within the European populations a north to south decline can be seen with Finnish (FIN) having the highest odds for matching the Archaic genome followed by British from England and Scotland (GBR) and Utah residents with ancestry from northern and western Europe (CEU). Toscani in Italy (TSI) and Iberians from Spain (IBS) show the lowest levels of matching (Figure 12).

**Figure 10:**
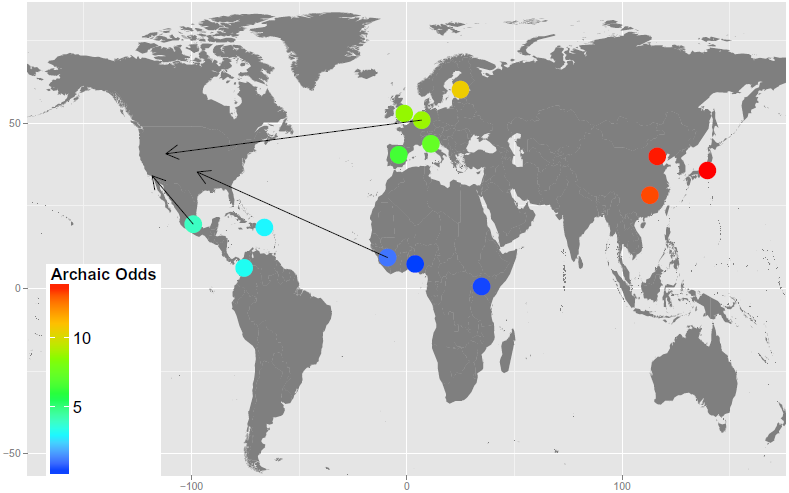
Odds scores of Fisher’s exact test for an enrichment of Archaic-matching IBD segments in different populations are represented by colored dots. The arrows point from the region the populations stem from to the region of sample collection. IBD segments that are shared by Asians match the Archaic genome significantly more often than IBD segments that are shared by other populations (red dots). Africans show the lowest matching with the Archaic genome (dark blue dots). Europeans show more matching than Admixed Americans (green and yellow vs. light blue dots).

**Figure 11:**
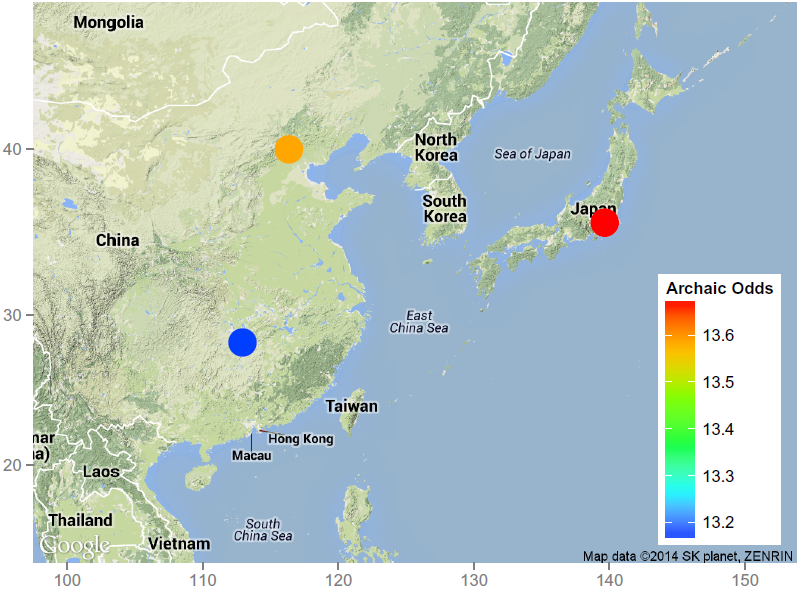
Odds scores of the Fisher’s exact tests for an enrichment of Archaic-matching IBD segments in Asian populations are represented by colored dots. Within the Asian populations Japanese (JPN) have slightly higher odds for matching the Archaic genome (red dot) than Han Chinese from South (CHS), while Han Chinese from Beijing (CHB) lie in between (blue and orange dots), but the differences are very small.

**Figure 12:**
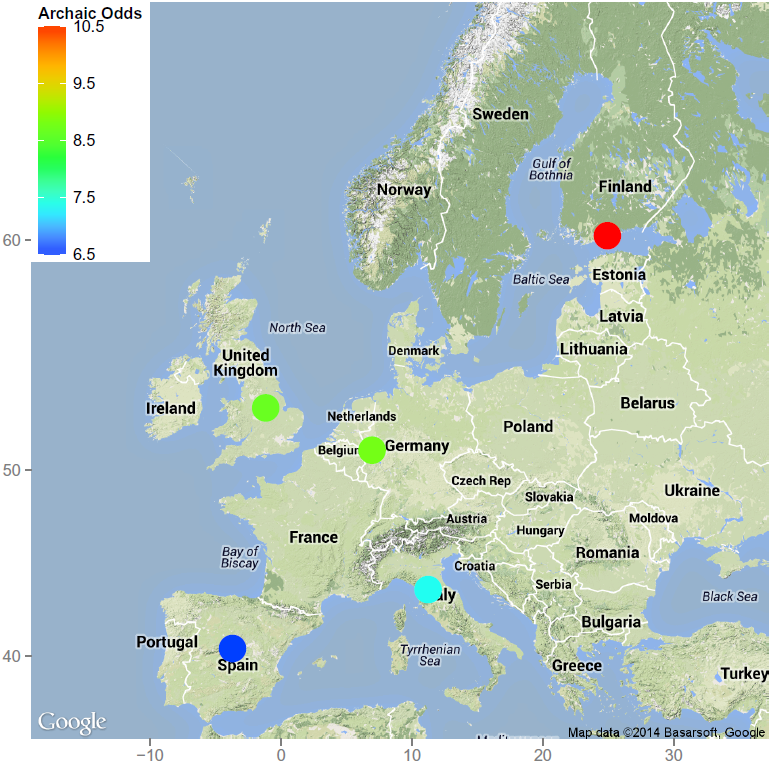
Odds scores of the Fisher’s exact tests for an enrichment of Archaic-matching IBD segments in European populations are represented by colored dots. Utah residents with ancestry from northern and western Europe (CEU) are symbolized by a dot in Central Europe. Within the European populations a north to south decline can be seen with Finnish (FIN) having the highest odds for matching the Archaic genome (red dot) followed by British from England and Scotland (GBR) and Utah residents with ancestry from northern and western Europe (CEU) (green dots). Toscani in Italy (TSI) and Iberians from Spain (IBS) show the lowest levels of matching (blue dots).

Using Fisher’s exact test on IBD segments of chromosome X, we found no clear enrichment of Denisova-matching segments in any of the tested populations and also very low differences in the odds for matching the Archaic genome. Thus, we confirmed our findings from the above correlation analysis. Again, results for chromosome X of the Neandertal genome are similar to those of the autosomes although the odds are generally lower. Figure 13 shows the odds scores of the Fisher’s exact tests for an enrichment of Neandertal-matching IBD segments of chromosome X in different populations. As expected, Asians again show the highest odds for IBD segments matching the Neandertal genome (odds ratio of 7.02 and *p*-value of 2.06e-143), while Africans have the lowest odds (odds ratio of 0.15 and *p*-value of 1.17e-151). Europeans show more matching with the Neandertal genome than Admixed Americans (odds ratio of 4.28 vs. 2.67). Within the Asian populations Japanese (JPN) have slightly higher odds for matching the Neandertal genome than Han Chinese from Beijing (CHB), while Han Chinese from South (CHS) lie in between, but the differences are again very small (Figure 14). Within the European populations a north to south decline can be seen with Finnish (FIN) having the highest odds for matching the Neandertal genome followed by British from England and Scotland (GBR) and Utah residents with ancestry from northern and western Europe (CEU). Toscani in Italy (TSI) and Iberians from Spain (IBS) show the lowest levels of matching (Figure 15).

**Figure 13:**
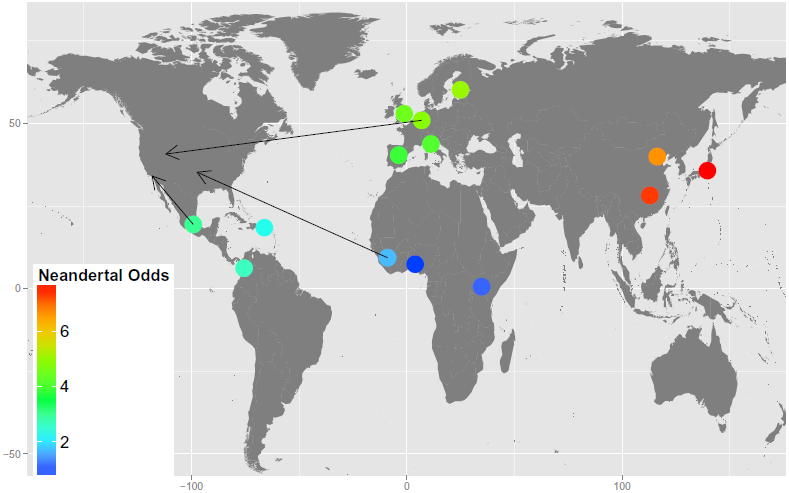
Odds scores of Fisher’s exact test for an enrichment of Neandertal-matching IBD segments on chromosome X in different populations are represented by colored dots. The arrows point from the region the populations stem from to the region of sample collection. IBD segments that are shared by Asians match the Neandertal genome significantly more often than IBD segments that are shared by other populations (red dots). Africans show the lowest matching with the Neandertal genome (dark blue dots). Europeans show more matching than Admixed Americans (green vs. light blue dots).

**Figure 14:**
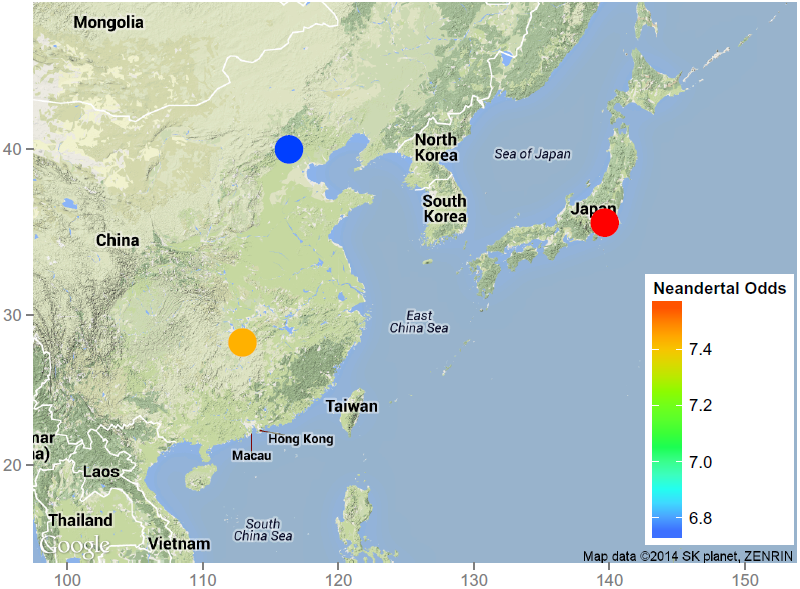
Odds scores of the Fisher’s exact tests for an enrichment of Neandertal-matching IBD segments on chromosome X in Asian populations are represented by colored dots. Within the Asian populations Japanese (JPN) have slightly higher odds for matching the Neandertal genome (red dot) than Han Chinese from Beijing (CHB), while Han Chinese from South (CHS) lie in between (blue and orange dots), but the differences are very small.

**Figure 15:**
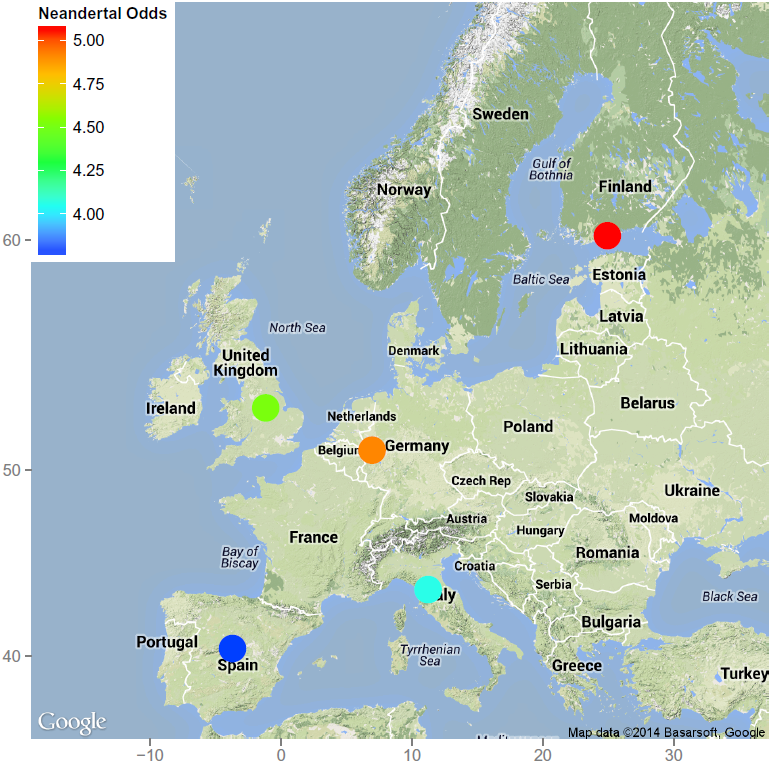
Odds scores of the Fisher’s exact tests for an enrichment of Neandertal-matching IBD segments on chromosome X in European populations are represented by colored dots. Utah residents with ancestry from northern and western Europe (CEU) are symbolized by a dot in Central Europe. Within the European populations a north to south decline can be seen with Finnish (FIN) having the highest odds for matching the Neandertal genome (red dot) followed by Utah residents with ancestry from northern and western Europe (CEU) and British from England and Scotland (GBR) (orange and green dots). Toscani in Italy (TSI) and Iberians from Spain (IBS) show the lowest levels of matching (blue dots).

In order to concentrate on strong effects in terms of population proportions, we investigated which population has the majority proportion for an IBD segment that matches a particular genome — the population to which the majority of the individuals possessing this segment belong to. Figure 16 shows the population with the majority proportion for each IBD segment. The IBD segments are presented for each genome, where the colors show the populations with majority proportion for the according IBD segment. For the autosomes, more than half of the Neandertal-matching IBD segments have Asians or Europeans as majority population proportions. However, still many Neandertal-matching segments are mainly shared by Africans which to some extend contradicts the hypothesis of Prüfer et al. (39) that Neandertal ancestry in Africans is due to back-to-Africa gene flow. For the Archaic genome (intersection of Neandertal-and Denisova-matching IBD segments), IBD segments dominated by Asians or Europeans are also enriched if compared to all IBD segments found in the 1000 Genomes Project data (we will call the set of these segments “human genome”). The enrichment by Asian or European IBD segments is lower for the Denisova genome, but still significant, especially for Asian segments. Furthermore, as can be seen in Figure 16, more IBD segments are found that match the Neandertal genome than segments that match the Denisova genome. One explanation for this is that the unknown ancient gene flow into Denisovans mentioned by Prüfer et al. (39) replaced some segments from the common ancestor of Neandertals and Denisovans in the Denisova genome. Therefore, these segments are only shared by modern humans and Neandertals although they might have been introduced by the aforementioned common ancestor. Another explanation is that ancestors of modern humans experienced more admixture events with Neandertals than with Denisovans. We assume that most Archaic-matching IBD segments found in Africans (IBD segments shared by all genomes) are old probably stemming from common ancestors of these hominid groups.

**Figure 16:**
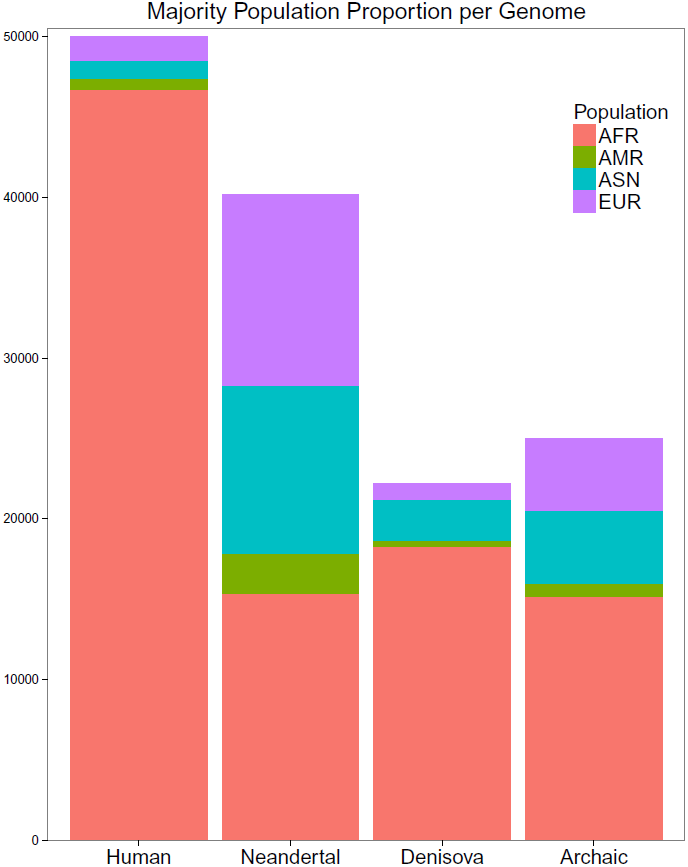
For each IBD segment, the population with the majority proportion is determined. IBD segments are given for each matching genome, where the color indicates the population that has majority proportion. For the human genome, 50,000 random IBD segments were chosen. More than half of the Neandertal-matching IBD segments have Asians or Europeans as majority population. The Archaic genome (Neandertal and Denisova) shows also an enrichment of IBD segments that are found mostly in Asians or Europeans. Denisova-matching IBD segments are often shared mainly by Asians.

Figure 17 shows the same results for chromosome X. IBD segments are presented for each genome, where the colors show the populations with majority proportion for the according IBD segment. For the X chromosome the differences are not as clear as for the autosomes, but still many of the Neandertal-matching IBD segments have Asians or Europeans as majority population proportions. There is hardly any enrichment by Asian or European IBD segments for the Denisova genome, in contrast to the results from the autosomes. Comparing the number of IBD segments across genomes, again more IBD segments are found on the X chromosome that match the Neandertal genome than segments that match the Denisova genome. In contrast to the results from the autosomes, even more IBD segments match the Archaic genome, that is, they match both the Neandertal and Denisova genome, than match the Neandertal genome. Most of the Archaic-matching IBD segments have Africans as the majority population, which is more prominent than in autosomes. Therefore, we conclude that most Archaic-matching IBD segments on the X chromosome are very old and, as on the autosomes, stem from common ancestors of the considered hominid groups. The relatively high number of Archaic-matching IBD segments dominated by Asians or Europeans might be explained by the lower diversity and therefore smaller divergence on the X chromosome (45). Neandertals may have introduced segments of DNA into the genomes of Europeans and Asians that still match the Denisova genome.

**Figure 17:**
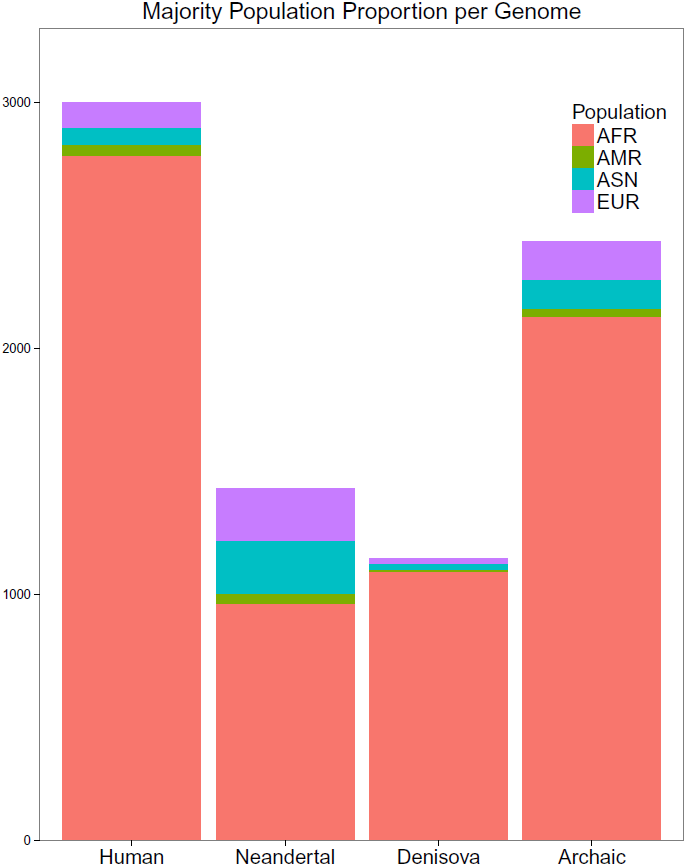
For each IBD segment of chromosome X, the population with the majority proportion is determined. IBD segments are given for each matching genome, where the color indicates the population that has majority proportion. For the human genome, 3,000 random IBD segments were chosen. Many of the Neandertal-matching IBD segments have Asians or Europeans as majority population. The Archaic genome (Neandertal and Denisova) shows also an enrichment of IBD segments that are found mostly in Asians or Europeans. Denisova-matching IBD segments are mostly shared by Africans.

#### 3.2.2 Densities of Population Proportions and Ancient Genomes

We plotted densities of population proportions for IBD segments that match a particular ancient genome (30% or more tagSNVs match) and for those that do not match that genome. Figure 18 shows the density of Asian proportions of Denisova-matching IBD segments (red) vs. the density of Asian proportions of non-Denisova-matching IBD segments (blue). Figure 19 shows analogous densities for Europeans. In comparison to all populations the density of Denisova-matching IBD segments that are observed in Asians and Europeans is higher than for non-matching IBD segments. This can be seen by the higher densities of matching IBD segments (red) compared to densities of non-matching IBD segments (blue) if the population proportions are not very close to zero — or, conversely, it can be seen at the lower peak at zero (less IBD segments that match the Denisova genome without Asian or European sharing). Many Denisova-matching IBD segments are shared exclusively among Asians which is indicated by the high density at a population proportion of 1 (red density in Figure 18). A population proportion of 1 means that IBD segments are exclusively shared among the respective population.

**Figure 18:**
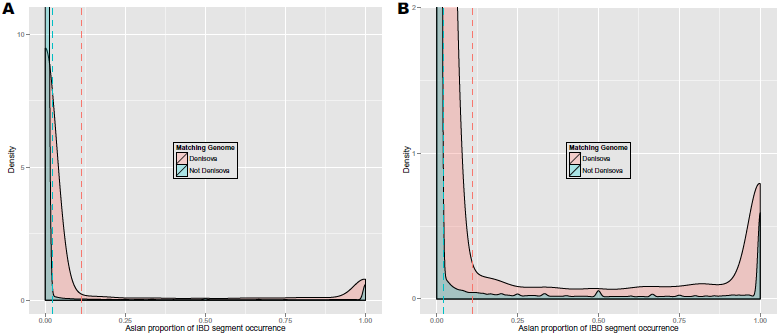
**Panel A**: Density of Asian proportions of Denisova-matching IBD segments (red) vs. density of Asian proportions of non-Denisova-matching IBD segments (blue). IBD segments were extracted from phased genotyping data of the 1000 Genomes Project. Dotted lines indicate the respective means. **Panel B**: The same densities as in Panel A but zoomed in.

**Figure 19:**
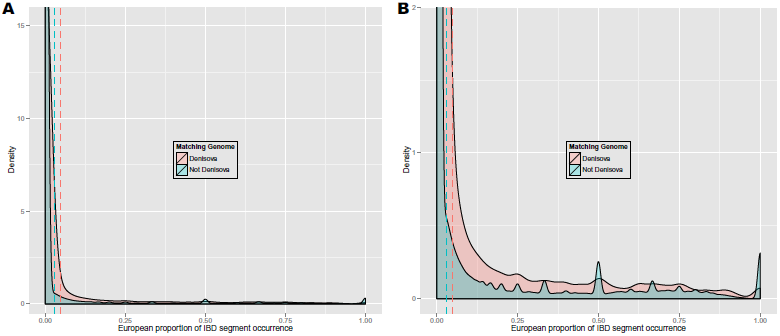
**Panel A**: Density of European proportions of Denisova-matching IBD segments (red) vs. density of European proportions of non-Denisova-matching IBD segments (blue). IBD segments were extracted from phased genotyping data of the 1000 Genomes Project. Dotted lines indicate the respective means. **Panel B**: The same densities as in Panel A but zoomed in.

**Figure 20:**
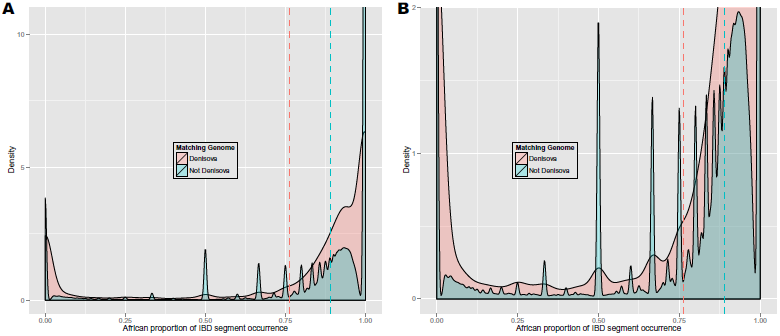
**Panel A**: Density of African proportions of Denisova-matching IBD segments (red) vs. density of African proportions of non-Denisova-matching IBD segments (blue). IBD segments were extracted from phased genotyping data of the 1000 Genomes Project. Dotted lines indicate the respective means. Peaks for non-Denisova-matching IBD segments are found at 0.5, 0.66, 0.33, 0.75, and 0.8, which corresponds to 1/2, 2/3, 1/3, 3/4, 4/5 (number of Africans / total number of individuals that have the IBD segment). The density of African proportions of Denisova-matching IBD segments has two peaks: one at a low and one at a high proportion of Africans. **Panel B**: The same densities as in Panel A but zoomed in.

Figures 21 and 22 show analogous densities as in Figures 18 and 19, but for the Neandertal genome. The differences we already observed for the Denisova genome are even more prominent for the Neandertal: Neandertal-matching IBD segments are observed even more often in Asians and Europeans than non-matching IBD segments if compared to all segments of the respective category. The higher densities (red) at a population proportion not close to zero are now more prominent — or conversely, the lower peak at zero for Neandertal-matching IBD segments becomes clearer (less IBD segments that match the Neandertal genome without Asian or European sharing). For Asians, the peak at 1 in Figure 21 is even higher, representing IBD segments that are shared exclusively among Asians. IBD segment sharing exclusively within Asians is rather common, as the blue peaks at 1 in Figure 18 and Figure 21 show.

**Figure 21:**
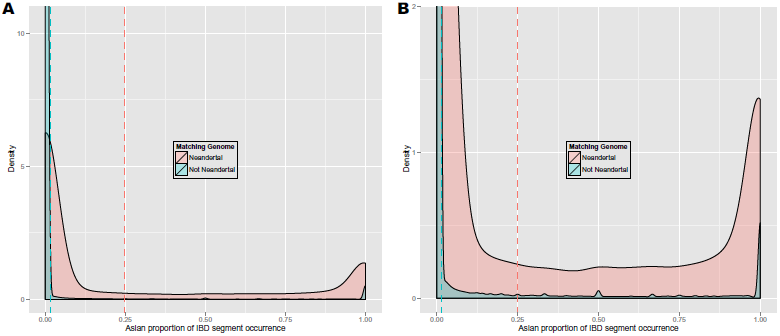
**Panel A**: Density of Asian proportions of Neandertal-matching IBD segments (red) vs. density of Asian proportions of non-Neandertal-matching IBD segments (blue). IBD segments were extracted from phased genotyping data of the 1000 Genomes Project. Dotted lines indicate the respective means. Many Neandertal-enriched IBD segments are shared mainly or exclusively by Asians as the peak at a proportion close to 1 shows. **Panel B**: The same densities as in Panel A but zoomed in.

**Figure 22:**
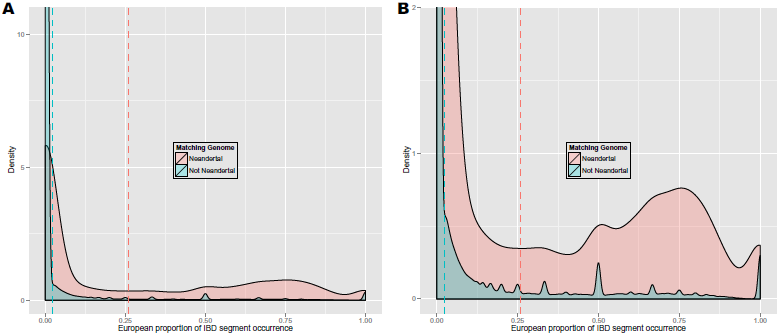
**Panel A**: Density of European proportions of Neandertal-matching IBD segments (red) vs. density of European proportions of non-Neandertal-matching IBD segments (blue). IBD segments were extracted from phased genotyping data of the 1000 Genomes Project. Dotted lines indicate the respective means. Many Neandertal-enriched IBD segments are shared mainly or exclusively by Asians as the peak at a proportion close to 1 in Figure 21 shows. Neandertal-enriched IBD segments are often shared by Europeans but fewer segments are shared exclusively among Europeans than among Asians. **Panel B**: the same density zoomed in.

Figures 20 and 23 show the same densities for Africans. Both the density of African proportions of Denisova-matching and the same density for Neandertal-matching IBD segments have two peaks: one at a low and one at a high proportion of Africans. For Neandertal-matching IBD segments, the density at low proportions of Africans is even larger than for high proportions. Thus, IBD segments that match ancient genomes are either shared by a very low or a very high proportion of Africans. The low proportion of African density peak hints at admixture of ancestors of modern humans and Denisovans / Neandertals outside of Africa or the back to Africa gene flow mentioned by Prüfer et al. (39). The density peak at high proportions of Africans may be due to ancient DNA segments shared by hominid groups that were lost in other continental populations. This finding could support the hypothesis that ancient population substructure in Africa also allowed for the occurrence of different continental patterns of DNA sharing between modern humans and ancient genomes (10, 11, 16).

**Figure 23:**
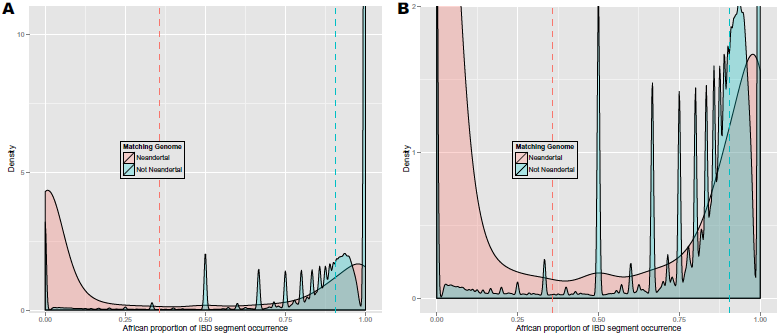
**Panel A**: Density of African proportions of Neandertal-matching IBD segments (red) vs. density of African proportions of non-Neandertal-matching IBD segments (blue). IBD segments were extracted from phased genotyping data of the 1000 Genomes Project. Dotted lines indicate the respective means. Peaks for non-Neandertal-matching IBD segments are found at 0.5, 0.66, 0.33, 0.75, and 0.8, which corresponds to 1/2, 2/3, 1/3, 3/4, 4/5 (number of Africans / total number of individuals that have the IBD segment). The density of African proportions of Neandertal-matching IBD segments has two peaks: one at a low and one at a high proportion of Africans. The density of low proportions of Africans is even larger than the density of high proportions. **Panel B**: the same density zoomed in.

IBD segments that match the “Archaic genome” are those IBD segments that match both the Denisova and Neandertal genome. Population proportion densities for the Archaic genome are presented in Figures 24, 25, and 26 for Asians, Europeans, and Africans, respectively. For the Ar-chaic genome, we see the same figure as for the Neandertal and the Denisova genome: the African density is bimodal that means either an IBD segment is dominated by Africans or it contains no or only few Africans.

**Figure 24:**
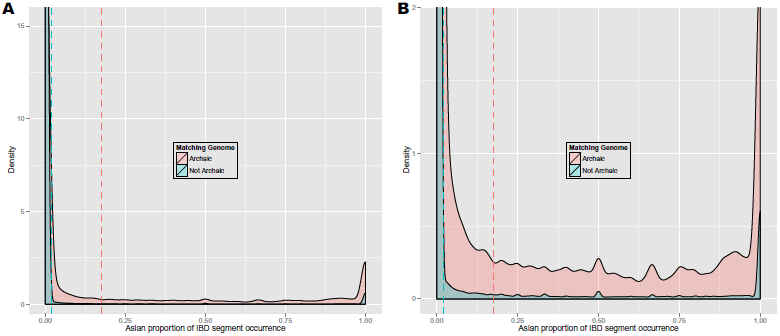
**Panel A**: Density of Asian proportions of Archaic-matching IBD segments (red) vs. density of Asian proportions of non-Archaic-matching IBD segments (blue). IBD segments were extracted from phased genotyping data of the 1000 Genomes Project. Dotted lines indicate the respective means. **Panel B**: the same density zoomed in.

**Figure 25:**
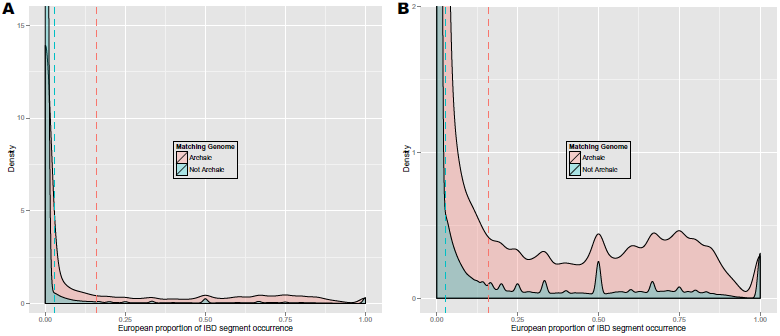
**Panel A**: Density of European proportions of Archaic-genome- matching IBD segments (red) vs. density of European proportions of non-Archaic-matching IBD segments (blue). IBD segments were extracted from phased genotyping data of the 1000 Genomes Project. Dotted lines indicate the respective means. Many Archaic-genome-enriched IBD segments are shared mainly or exclusively by Asians as the peak at a proportion close to 1 shows. **Panel B**: the same densities zoomed in.

**Figure 26:**
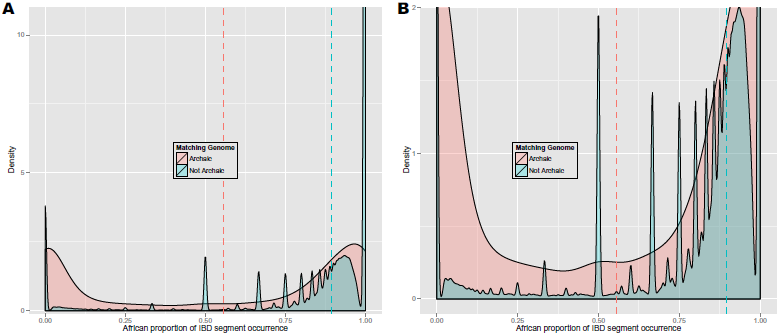
**Panel A**: Density of African proportions of Archaic-matching IBD segments (red) vs. density of African proportions of non-Archaic- matching IBD segments (blue). IBD segments were extracted from phased genotyping data of the 1000 Genomes Project. Dotted lines indicate the respective means. Peaks for non-Archaic-matching IBD segments are found at 0.5, 0.66, 0.33, 0.75, and 0.8, which corresponds to 1/2, 2/3, 1/3, 3/4, 4/5 (number of Africans / total number of individuals that have the IBD segment). The density of African proportions of Archaic-matching IBD segments has two peaks: one at a low and one at a high proportion of Africans. **Panel B**: the same density zoomed in.

Figures 27–35 show the same densities for chromosome X. Again, we could confirm our novel finding, that Denisova- and Archaic-matching segments on chromosome X are mainly shared by Africans. Only very low proportions of Asians or Europeans share Denisova- or Archaic-matching IBD segments. For Neandertal-matching IBD segments, the density is higher near an African proportion of 1 compared to the autosomes. However, also Asians have a high density peak at a population proportion close to 1 and also Europeans show a small peak. Both peaks are clearly higher than the respective peaks for non-matching IBD segments.

**Figure 27:**
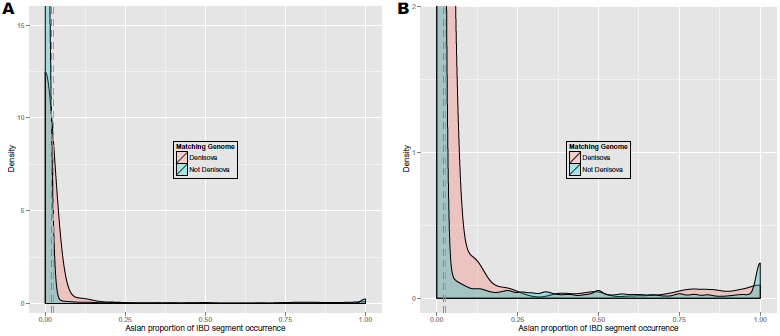
**Panel A**: Density of Asian proportions of Denisova-matching IBD segments on chromosome X (red) vs. density of Asian proportions of non-Denisova-matching IBD segments (blue). IBD segments were extracted from phased genotyping data of the 1000 Genomes Project. Dotted lines indicate the respective means. **Panel B**: The same densities as in Panel A but zoomed in.

**Figure 28:**
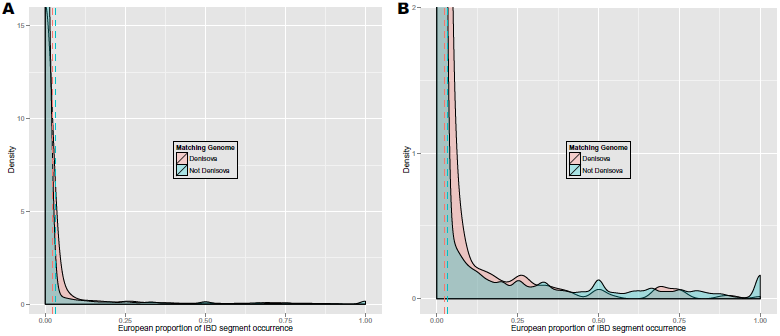
**Panel A**: Density of European proportions of Denisova- matching IBD segments on chromosome X (red) vs. density of European proportions of non-Denisova-matching IBD segments (blue). IBD segments were extracted from phased genotyping data of the 1000 Genomes Project. Dotted lines indicate the respective means. **Panel B**: The same densities as in Panel A but zoomed in.

**Figure 29:**
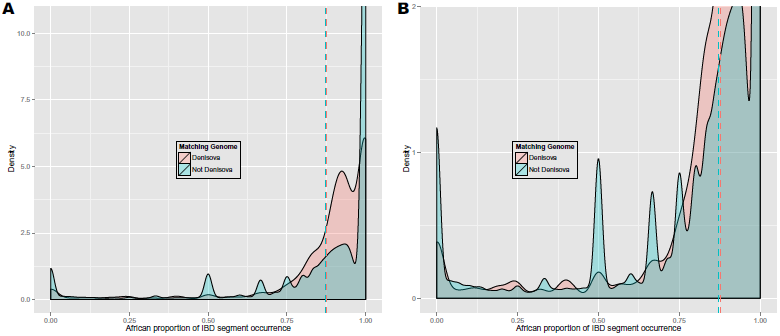
**Panel A**: Density of African proportions of Denisova-matching IBD segments on chromosome X (red) vs. density of African proportions of non-Denisova-matching IBD segments (blue). IBD segments were extracted from phased genotyping data of the 1000 Genomes Project. Dotted lines indicate the respective means. More segments are shared by many Africans than by only a few or no African which is indicated by the higher densities at high African proportion compared to low African proportion both for Denisova-matching and non-Denisova-matching IBD segments. **Panel B**: The same densities as in Panel A but zoomed in.

**Figure 30:**
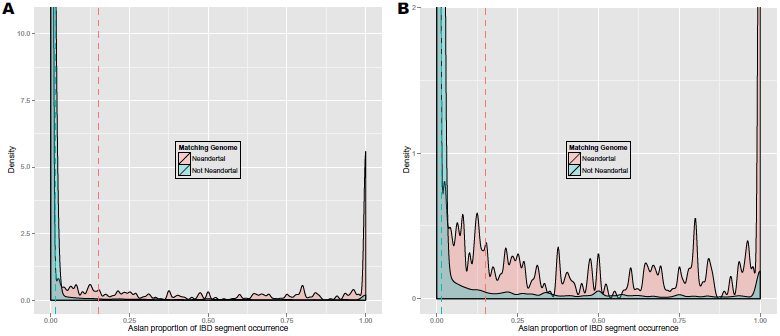
**Panel A**: Density of Asian proportions of Neandertal-matching IBD segments on chromosome X (red) vs. density of Asian proportions of non-Neandertal-matching IBD segments (blue). IBD segments were extracted from phased genotyping data of the 1000 Genomes Project. Dotted lines indicate the respective means. Many Neandertal-enriched IBD segments are shared mainly or exclusively by Asians as the peak at a proportion close to 1 shows. **Panel B**: The same densities as in Panel A but zoomed in.

**Figure 31:**
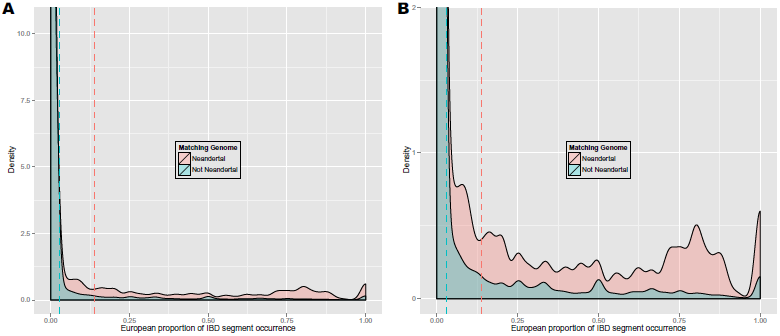
**Panel A**: Density of European proportions of Neandertal-matching IBD segments on chromosome X (red) vs. density of European proportions of non-Neandertal-matching IBD segments (blue). IBD segments were extracted from phased genotyping data of the 1000 Genomes Project. Dotted lines indicate the respective means. Many Neandertal-enriched IBD segments are shared mainly or exclusively by Asians as the peak at a proportion close to 1 in Figure 30 shows. Neandertal-enriched IBD segments are often shared by Europeans but fewer segments are shared exclusively among Europeans than among Asians. **Panel B**: the same densities zoomed in.

**Figure 32:**
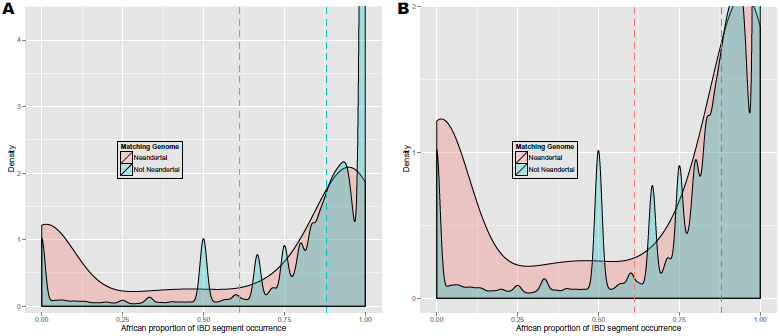
**Panel A**: Density of African proportions of Neandertal-matching IBD segments on chromosome X (red) vs. density of African proportions of non-Neandertal-matching IBD segments (blue). IBD segments were extracted from phased genotyping data of the 1000 Genomes Project. Dotted lines indicate the respective means. More segments are shared by many Africans than by only a few or no African which is indicated by the higher densities at high African proportion compared to low African proportion both for Neandertal-matching and non-Neandertal-matching IBD segments. **Panel B**: the same densities zoomed in.

**Figure 33:**
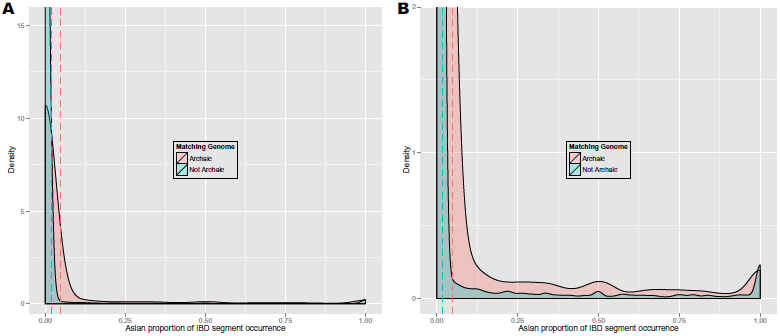
**Panel A**: Density of Asian proportions of Archaic-matching IBD segments on chromosome X (red) vs. density of Asian proportions of non-Archaic-matching IBD segments (blue). IBD segments were extracted from phased genotyping data of the 1000 Genomes Project. Dotted lines indicate the respective means. **Panel B**: the same densities zoomed in.

**Figure 34:**
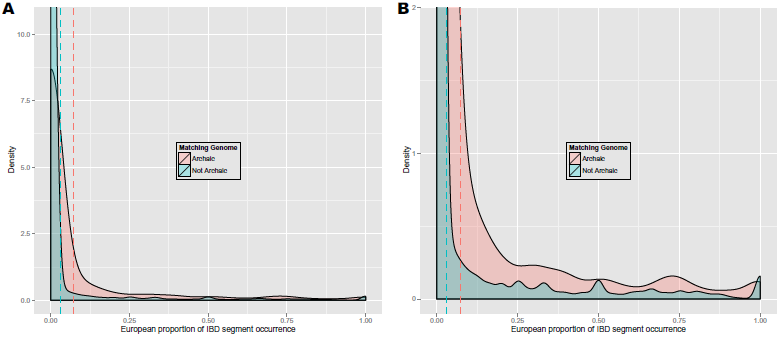
**Panel A**: Density of European proportions of Archaic-matching IBD segments on chromosome X (red) vs. density of European proportions of non-Archaic-matching IBD segments (blue). IBD segments were extracted from phased genotyping data of the 1000 Genomes Project. Dotted lines indicate the respective means. **Panel B**: the same densities zoomed in.

**Figure 35:**
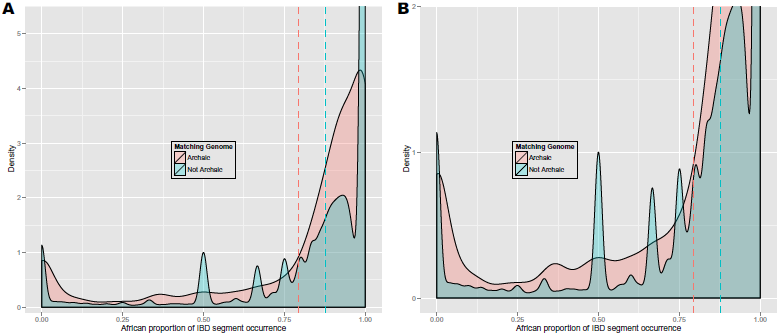
**Panel A**: Density of African proportions of Archaic-matching IBD segments on chromosome X (red) vs. density of African proportions of non-Archaic-matching IBD segments (blue). IBD segments were extracted from phased genotyping data of the 1000 Genomes Project. Dotted lines indicate the respective means. More segments are shared by many Africans than by only a few or no African which is indicated by the higher densities at high African proportion compared to low African proportion both for Archaic-matching and non-Archaic-matching IBD segments. **Panel B**: the same densities zoomed in.

#### 3.2.3 IBD Sharing Between Human Populations and Ancient Genomes on Different Chromosomes

The amount of IBD sharing with ancient genomes is not evenly distributed over all chromosomes. Table 3 lists the number of IBD segments that are exclusively shared by a certain continental population for each chromosome. Overall, it can be seen, that Europeans and Admixed Americans share fewer IBD segments exclusively than Asians. One reason for this is, that many IBD segments that are shared by Europeans can also be found in Admixed Americans and vice versa. Table 4 shows for each chromosome the number of Denisova-matching IBD segments that are shared exclusively by different continental populations. Tables 5 and 6 show the same information for Neandertal- and Archaic-matching IBD segments, respectively. Europeans and Admixed Americans share hardly any IBD segments exclusively with the Denisova in contrast to Asians and Africans. Africans share a smaller fraction of IBD segments with the Denisova than Asians if compared to the total number of exclusively shared segments (see Table 3). Considerably more Neandertal-matching than Denisova-matching IBD segments are exclusively shared by Europeans, but also Asians show even more segments matching the Neandertal genome than matching the Denisova. For Africans the number of Neandertal- and Denisova-matching IBD sgements are in a similar range. Counts for IBD segments matching the Archaic genome are generally lower than for Neandertals, but comparable to the Denisova. An exception is the number of Archaic-matching IBD segments found exclusively on the X chromosome of Africans which is considerably higher.

**Table 3:** Number of IBD segments shared exclusively by a particular continental population for each chromosome.

| Chr. | All | ASN | EUR | AMR | AFR |
| --- | --- | --- | --- | --- | --- |
| 1 | 105,615 | 1,418 | 557 | 365 | 65,131 |
| 2 | 115,235 | 1,243 | 706 | 382 | 71,449 |
| 3 | 94,257 | 984 | 471 | 344 | 58,488 |
| 4 | 95,029 | 1,180 | 492 | 326 | 59,566 |
| 5 | 92,342 | 1,149 | 555 | 451 | 56,478 |
| 6 | 83,432 | 1,293 | 520 | 313 | 51,691 |
| 7 | 79,543 | 708 | 424 | 388 | 49,027 |
| 8 | 77,934 | 625 | 409 | 383 | 48,546 |
| 9 | 59,377 | 627 | 336 | 225 | 37,772 |
| 10 | 64,694 | 884 | 392 | 308 | 38,326 |
| 11 | 66,613 | 673 | 432 | 250 | 40,787 |
| 12 | 63,508 | 773 | 328 | 199 | 39,889 |
| 13 | 48,327 | 608 | 309 | 205 | 29,408 |
| 14 | 44,645 | 764 | 258 | 133 | 27,185 |
| 15 | 40,663 | 522 | 198 | 162 | 25,957 |
| 16 | 42,915 | 260 | 306 | 162 | 27,687 |
| 17 | 37,823 | 349 | 196 | 186 | 25,041 |
| 18 | 39,027 | 392 | 169 | 145 | 24,181 |
| 19 | 29,580 | 311 | 165 | 179 | 18,457 |
| 20 | 30,314 | 235 | 154 | 150 | 18,855 |
| 21 | 19,108 | 187 | 112 | 78 | 11,590 |
| 22 | 18,339 | 253 | 132 | 72 | 12,258 |
| X | 43,069 | 335 | 251 | 183 | 23,087 |

The column labeled “Chr.” gives the chromosome, “All” reports the total number of IBD segment per chromosome, “ASN”, “EUR”, “AMR”, and “AFR” report the number of IBD segments shared exclusively by Asians, Europeans, Admixed Americans, and Africans, respectively.

**Table 4:** Number of Denisova-matching IBD segments shared exclusively by a particular continental population for each chromosome.

| Chr. | Denisova | Denisova-ASN | Denisova-EUR | Denisova-AMR | Denisova-AFR |
| --- | --- | --- | --- | --- | --- |
| 1 | 1,908 | 182 | 13 | 5 | 522 |
| 2 | 1,825 | 134 | 5 | 2 | 542 |
| 3 | 1,387 | 36 | 3 | 2 | 479 |
| 4 | 1,796 | 174 | 0 | 17 | 561 |
| 5 | 1,552 | 184 | 5 | 1 | 412 |
| 6 | 1,511 | 101 | 2 | 0 | 473 |
| 7 | 1,287 | 80 | 2 | 11 | 366 |
| 8 | 1,194 | 41 | 2 | 0 | 389 |
| 9 | 1,018 | 29 | 3 | 1 | 368 |
| 10 | 1,141 | 76 | 4 | 8 | 336 |
| 11 | 1,064 | 73 | 18 | 8 | 332 |
| 12 | 876 | 54 | 0 | 1 | 287 |
| 13 | 791 | 81 | 5 | 0 | 221 |
| 14 | 682 | 36 | 0 | 4 | 169 |
| 15 | 747 | 48 | 2 | 0 | 284 |
| 16 | 647 | 10 | 3 | 4 | 245 |
| 17 | 508 | 36 | 1 | 5 | 177 |
| 18 | 591 | 31 | 0 | 0 | 203 |
| 19 | 534 | 22 | 2 | 1 | 142 |
| 20 | 444 | 14 | 0 | 1 | 121 |
| 21 | 373 | 20 | 0 | 0 | 97 |
| 22 | 310 | 25 | 5 | 0 | 100 |
| X | 1,147 | 7 | 1 | 1 | 351 |

The column labeled “Chr.” gives the chromosome, “Denisova” reports the number of Denisova-matching IBD segments, “Denisova-ASN”, “Denisova-EUR”, “Denisova-AMR”, and “Denisova-AFR” report the number of Denisova-matching IBD segments shared exclusively by Asians, Europeans, Admixed Americans, and Africans, respectively.

**Table 5:** Number of Neandertal-matching IBD segments shared exclusively by a particular continental population for each chromosome.

| Chr. | Neandertal | Neandertal-ASN | Neandertal-EUR | Neandertal-AMR | Neandertal-AFR |
| --- | --- | --- | --- | --- | --- |
| 1 | 3,266 | 464 | 83 | 11 | 392 |
| 2 | 3,329 | 385 | 146 | 31 | 472 |
| 3 | 2,788 | 351 | 90 | 11 | 377 |
| 4 | 2,962 | 316 | 79 | 19 | 491 |
| 5 | 2,691 | 404 | 72 | 31 | 347 |
| 6 | 2,784 | 372 | 77 | 19 | 350 |
| 7 | 2,275 | 212 | 65 | 20 | 332 |
| 8 | 2,034 | 186 | 55 | 19 | 373 |
| 9 | 1,831 | 221 | 69 | 6 | 258 |
| 10 | 1,916 | 313 | 72 | 16 | 168 |
| 11 | 2,133 | 203 | 80 | 11 | 232 |
| 12 | 1,994 | 267 | 47 | 6 | 200 |
| 13 | 1,536 | 206 | 42 | 18 | 203 |
| 14 | 1,435 | 303 | 48 | 6 | 119 |
| 15 | 1,217 | 187 | 26 | 13 | 152 |
| 16 | 1,026 | 69 | 45 | 8 | 178 |
| 17 | 924 | 102 | 36 | 15 | 151 |
| 18 | 1,204 | 131 | 38 | 11 | 193 |
| 19 | 869 | 120 | 20 | 15 | 112 |
| 20 | 935 | 76 | 40 | 5 | 145 |
| 21 | 526 | 52 | 28 | 1 | 90 |
| 22 | 514 | 69 | 26 | 6 | 70 |
| X | 1,429 | 108 | 34 | 3 | 328 |

The column labeled “Chr.” gives the chromosome, “Neandertal” reports the number of Neandertal-matching IBD segments, “Neandertal-ASN”, “Neandertal-EUR”, “Neandertal-AMR”, and “Neandertal-AFR” report the number of Neandertal-matching IBD segments shared exclusively by Asians, Europeans, Admixed Americans, and Africans, respectively.

**Table 6:** Number of Archaic-matching IBD segments shared exclusively by a particular continental population for each chromosome.

| Chr. | Archaic | Archaic-ASN | Archaic-EUR | Archaic-AMR | Archaic-AFR |
| --- | --- | --- | --- | --- | --- |
| 1 | 1,992 | 171 | 20 | 3 | 320 |
| 2 | 2,400 | 115 | 28 | 9 | 471 |
| 3 | 1,614 | 122 | 13 | 4 | 257 |
| 4 | 1,910 | 117 | 14 | 6 | 455 |
| 5 | 1,603 | 160 | 18 | 7 | 223 |
| 6 | 1,741 | 194 | 45 | 8 | 296 |
| 7 | 1,418 | 76 | 16 | 13 | 251 |
| 8 | 1,468 | 50 | 12 | 6 | 393 |
| 9 | 979 | 80 | 20 | 4 | 186 |
| 10 | 1,257 | 95 | 13 | 11 | 203 |
| 11 | 1,265 | 67 | 39 | 0 | 216 |
| 12 | 1,000 | 101 | 2 | 1 | 118 |
| 13 | 872 | 62 | 18 | 0 | 197 |
| 14 | 896 | 113 | 14 | 4 | 102 |
| 15 | 700 | 55 | 4 | 1 | 116 |
| 16 | 628 | 23 | 6 | 0 | 136 |
| 17 | 622 | 58 | 10 | 2 | 167 |
| 18 | 732 | 43 | 3 | 0 | 110 |
| 19 | 644 | 34 | 10 | 3 | 103 |
| 20 | 617 | 26 | 6 | 0 | 146 |
| 21 | 314 | 20 | 6 | 0 | 51 |
| 22 | 337 | 25 | 5 | 4 | 59 |
| X | 2,435 | 38 | 21 | 3 | 635 |

The column labeled “Chr.” gives the chromosome, “Archaic” reports the number of Archaic-matching IBD segments, “Archaic-ASN”, “Archaic-EUR”, “Archaic-AMR”, and “Archaic-AFR” report the number of Archaic-matching IBD segments shared exclusively by Asians, Europeans, Admixed Americans, and Africans, respectively.

Table 7 lists the number of IBD segments that are shared by at least one individual of a certain continental population for each chromosome. It can be seen, that Asians share fewer IBD segments non-exclusively than the other continental populations. Many IBD segments are shared by at least one Admixed American due to their admixed ancestry. Table 8 reports for each chromosome the number of Denisova-matching IBD segments that are shared by at least one individual of a certain continental population. Tables 9 and 10 show the same information for Neandertal- and Archaic-matching IBD segments, respectively. Europeans have numbers of Denisova-matching IBD segments that are similar to the numbers for Asians. Admixed Americans share clearly more of these segments. However, Asians still have the highest amount of Denisova-sharing if compared to the total number of segments shared by at least one individual of a certain population (see Table 7). For Neandertal-matching IBD segments the situation is similar, but surprisingly few of these segments are shared by at least one African. Archaic-matching IBD segments are especially often found on the X chromosome and are most often shared by Admixed Americans and Africans.

**Table 7:**
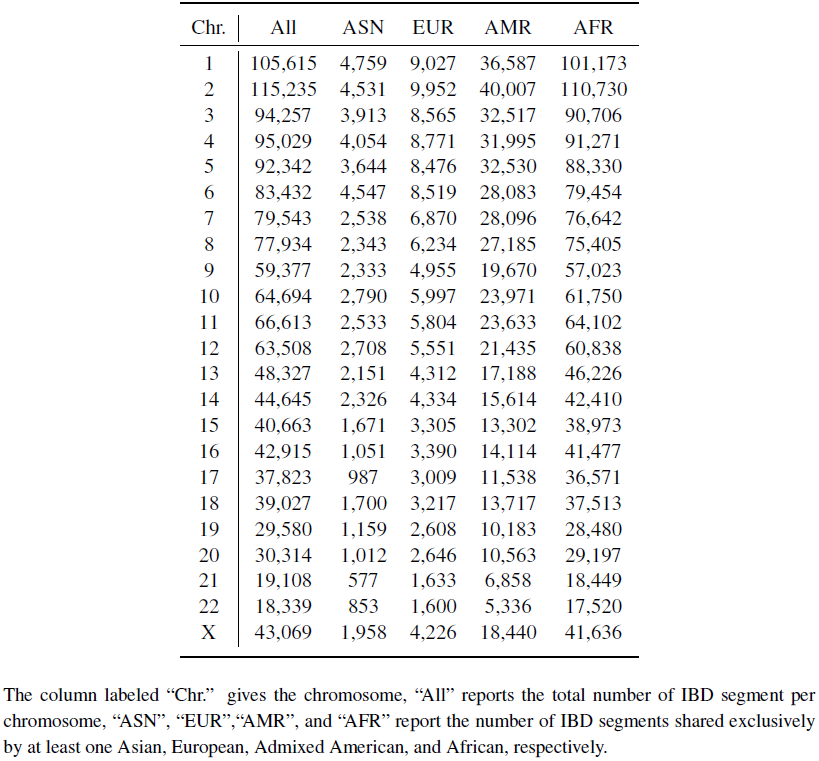
Number of IBD segments shared by at least one individual of a particular continental population for each chromosome.

**Table 8:** Number of Denisova-matching IBD segments shared by at least one individual of a particular continental population for each chromosome.

| Chr. | Denisova | Denisova-ASN | Denisova-EUR | Denisova-AMR | Denisova-AFR |
| --- | --- | --- | --- | --- | --- |
| 1 | 1,908 | 398 | 338 | 1,096 | 1,606 |
| 2 | 1,825 | 374 | 364 | 1,057 | 1,574 |
| 3 | 1,387 | 180 | 277 | 798 | 1,280 |
| 4 | 1,796 | 416 | 326 | 983 | 1,518 |
| 5 | 1,552 | 387 | 253 | 868 | 1,239 |
| 6 | 1,511 | 322 | 305 | 844 | 1,303 |
| 7 | 1,287 | 172 | 175 | 784 | 1,147 |
| 8 | 1,194 | 122 | 202 | 741 | 1,120 |
| 9 | 1,018 | 105 | 171 | 588 | 949 |
| 10 | 1,141 | 158 | 211 | 697 | 1,000 |
| 11 | 1,064 | 201 | 197 | 592 | 903 |
| 12 | 876 | 140 | 126 | 511 | 788 |
| 13 | 791 | 169 | 139 | 452 | 663 |
| 14 | 682 | 109 | 109 | 449 | 619 |
| 15 | 747 | 110 | 123 | 390 | 660 |
| 16 | 647 | 50 | 99 | 365 | 602 |
| 17 | 508 | 83 | 86 | 271 | 449 |
| 18 | 591 | 98 | 124 | 316 | 531 |
| 19 | 534 | 84 | 116 | 343 | 498 |
| 20 | 444 | 37 | 68 | 296 | 409 |
| 21 | 373 | 54 | 75 | 230 | 335 |
| 22 | 310 | 76 | 66 | 162 | 258 |
| X | 1,147 | 76 | 146 | 757 | 1,124 |

The column labeled “Chr.” gives the chromosome, “Denisova” reports the number of Denisova-matching IBD segments, “Denisova-ASN”, “Denisova-EUR”, “Denisova-AMR”, and “Denisova-AFR” report the number of Denisova-matching IBD segments shared by at least one Asian, European, Admixed American, and African, respectively.

**Table 9:** Number of Neandertal-matching IBD segments shared by at least one individual of a particular continental population for each.

| Chr. | Neandertal | Neandertal-ASN | Neandertal-EUR | Neandertal-AMR | Neandertal-AFR |
| --- | --- | --- | --- | --- | --- |
| 1 | 3,266 | 1,531 | 1,628 | 2,012 | 1,818 |
| 2 | 3,329 | 1,268 | 1,634 | 2,082 | 1,979 |
| 3 | 2,788 | 1,202 | 1,396 | 1,715 | 1,580 |
| 4 | 2,962 | 1,027 | 1,461 | 1,868 | 1,835 |
| 5 | 2,691 | 1,063 | 1,451 | 1,603 | 1,501 |
| 6 | 2,784 | 1,206 | 1,469 | 1,729 | 1,607 |
| 7 | 2,275 | 703 | 1,142 | 1,487 | 1,419 |
| 8 | 2,034 | 538 | 843 | 1,318 | 1,357 |
| 9 | 1,831 | 784 | 951 | 1,116 | 1,003 |
| 10 | 1,916 | 839 | 1,015 | 1,207 | 1,000 |
| 11 | 2,133 | 798 | 1,200 | 1,436 | 1,353 |
| 12 | 1,994 | 919 | 1,181 | 1,350 | 967 |
| 13 | 1,536 | 601 | 765 | 950 | 900 |
| 14 | 1,435 | 735 | 728 | 848 | 627 |
| 15 | 1,217 | 560 | 622 | 770 | 662 |
| 16 | 1,026 | 242 | 509 | 678 | 646 |
| 17 | 924 | 244 | 391 | 572 | 574 |
| 18 | 1,204 | 516 | 582 | 746 | 722 |
| 19 | 869 | 347 | 406 | 562 | 533 |
| 20 | 935 | 286 | 504 | 629 | 553 |
| 21 | 526 | 131 | 271 | 331 | 334 |
| 22 | 514 | 248 | 264 | 299 | 300 |
| X | 1,429 | 376 | 453 | 862 | 1,135 |

The column labeled “Chr.” gives the chromosome, “Neandertal” reports the number of Neandertal-matching IBD segments, “Neandertal-ASN”, “Neandertal-EUR”, “Neandertal-AMR”, and “Neandertal-AFR” report the number of Neandertal-matching IBD segments shared by at least one Asian, European, Admixed American, and African, respectively.

**Table 10:** Number of Archaic-matching IBD segments shared by at least one individual of a particular continental population for each.

| Chr. | Archaic | Archaic-ASN | Archaic-EUR | Archaic-AMR | Archaic-AFR |
| --- | --- | --- | --- | --- | --- |
| 1 | 1,992 | 870 | 891 | 1,234 | 1,422 |
| 2 | 2,400 | 728 | 951 | 1,546 | 1,920 |
| 3 | 1,614 | 606 | 658 | 1,090 | 1,281 |
| 4 | 1,910 | 614 | 601 | 1,139 | 1,563 |
| 5 | 1,603 | 526 | 716 | 1,031 | 1,177 |
| 6 | 1,741 | 675 | 720 | 1,009 | 1,231 |
| 7 | 1,418 | 351 | 492 | 972 | 1,127 |
| 8 | 1,468 | 238 | 406 | 958 | 1,253 |
| 9 | 979 | 274 | 369 | 624 | 746 |
| 10 | 1,257 | 439 | 557 | 856 | 951 |
| 11 | 1,265 | 353 | 620 | 833 | 997 |
| 12 | 1,000 | 431 | 436 | 686 | 732 |
| 13 | 872 | 285 | 368 | 537 | 664 |
| 14 | 896 | 476 | 416 | 585 | 617 |
| 15 | 700 | 265 | 294 | 451 | 522 |
| 16 | 628 | 126 | 230 | 435 | 527 |
| 17 | 622 | 131 | 173 | 351 | 498 |
| 18 | 732 | 274 | 270 | 542 | 586 |
| 19 | 644 | 187 | 237 | 427 | 516 |
| 20 | 617 | 117 | 206 | 417 | 513 |
| 21 | 314 | 61 | 109 | 210 | 250 |
| 22 | 337 | 125 | 146 | 221 | 248 |
| X | 2,435 | 314 | 600 | 1,607 | 2,289 |

The column labeled “Chr.” gives the chromosome, “Archaic” reports the number of Archaic-matching IBD segments, “Archaic-ASN”, “Archaic-EUR”, “Archaic-AMR”, and “Archaic-AFR” report the number of Archaic-matching IBD segments shared by at least one Asian, European, Admixed American, and African, respectively.

Figures 36–38 show the distribution of Denisova-, Neandertal-, and Archaic-matching IBD segments along the genome, where Africans are contrasted to non-Africans. Interestingly, while African shared Denisova-matching IBD segments can be found in nearly every region that is covered by enough sequencing reads, there are many DNA regions that lack non-African Denisova-matching segments. There is only a relatively small number of non-African Denisova-matching IBD segments, therefore they cannot span the whole genome. Nevertheless, these segments tend to accumulate in certain regions of the genome while large regions contain no Denisova-matching IBD segments that are only shared by non-Africans. For Neandertal-matching IBD segments there are also regions without any non-African segments, but except for the X chromosome there are approximately as many segments as for Africans. In concordance with Sankararaman et al. (43), we found many regions on the X chromosome where non-African Neandertal-matching IBD segments were absent. For segments matching the Archaic genome a large number of IBD segments on chromosome X are found in Africans whereas on the other chromosomes only slightly more African IBD segments can be found compared to non-African.

**Figure 36:**
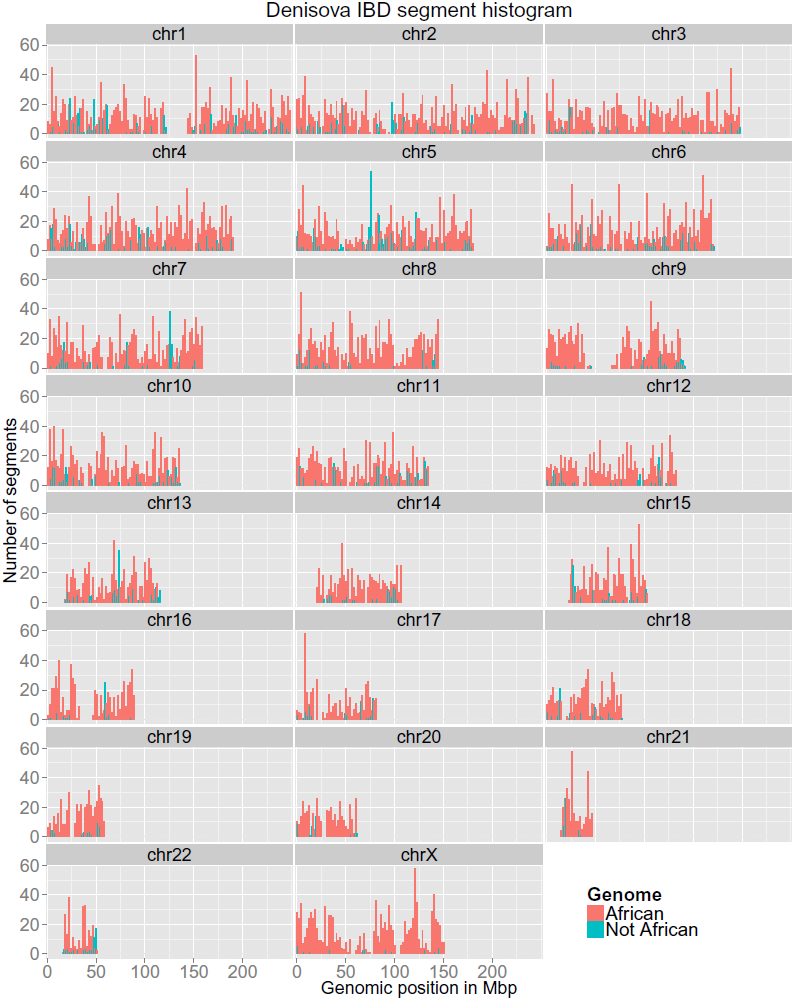
Histogram of Denisova-matching IBD segments along the genome. Segments shared by Africans are colored in red, segments shared by no African are colored in blue.

**Figure 37:**
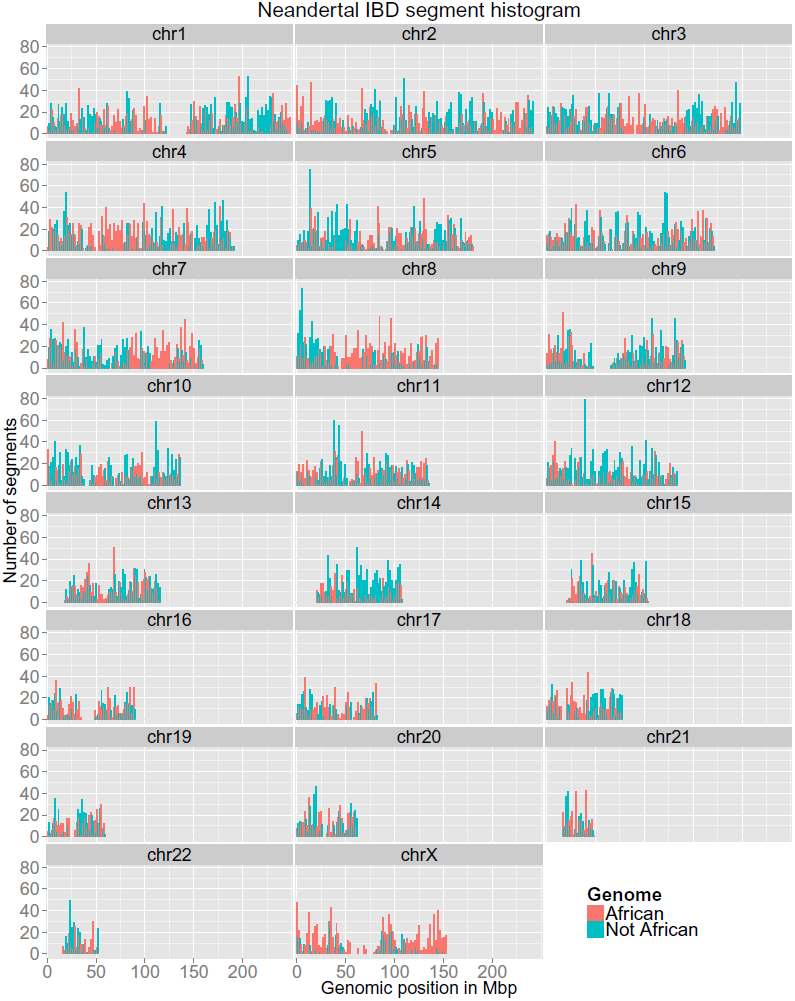
Histogram of Neandertal-matching IBD segments along the genome. Segments shared by Africans are colored in red, segments shared by no African are colored in blue.

**Figure 38:**
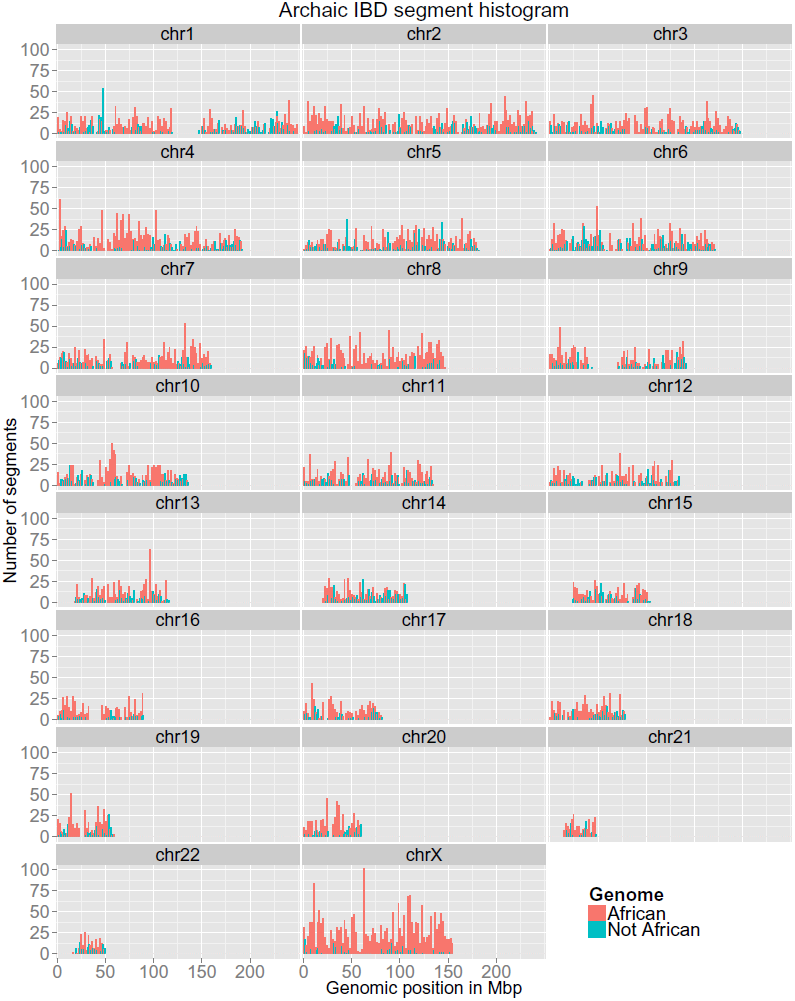
Histogram of Archaic-matching IBD segments along the genome. Segments shared by Africans are colored in red, segments shared by no African are colored in blue.

## 4 Analyses of Lengths of IBD Segments

### 4.1 Relating the IBD Length to Years from Present

We aimed to establish a relation between the length of an IBD segment and the time of the most recent common ancestor of the individuals that possess the IBD segment. The shorter the IBD segment is, the older it is assumed to be, the further in the past the most recent common ancestor should be found. For IBD length distributions, mathematical models have already been established. However, these models assume IBD segment sharing between only two individuals.

#### 4.1.1 Exponentially Distributed IBD Lengths

The length of an IBD segment is exponentially distributed with a mean of 100*/*(2*g*) cM (centi-Morgans), where *g* is the number of generations which separate the two haplotypes that share a segment from their common ancestor (4, 17, 38, 55, 56). Ulgen and Li (57) recommend to use a recombination rate, cM-to-Mbp ratio, of 1, however it varies from 0 to 9 along a chromosome (62).

For chromosome X, the sex-averaged recombination rate is 2*/*3 of the female recombination rate on the X chromosome (20). The female recombination rate on the X chromosome has approximately the same value as the sex-averaged autosomal recombination rate (26). Thus, we assume that the sex-averaged recombination rate on the X chromosome is 2*/*3 times the autosomal recombination rate.

**We are not able to perform reliable age estimations of the IBD segments based on their length.**

We encountered severe problems in estimating the age of IBD segments based on their length:

- The original ancestor DNA sequence is assumed to have a length of 1 Morgan before it is broken up by recombinations. However, founder genomes cannot be assumed to be distinguishable across the length of 1 Morgan.
- It is assumed that recombinations are random and all resulting segments have the same chance to survive. However, e.g. after population admixture or introgression of ancient genomes into ancestors of humans, recombined segments may have different fitness and some may vanish due to the high selective pressure. Thus, after such events the selective pressure leads to a bias of the IBD length distribution which makes the estimation of their age intractable.
- The age estimations are based on the mean, thus it is assumed that there are enough recombination events in each generation to average out random effects. Therefore, for few admixture/introgression events (few matings/offspring) these estimations are not reliable.

Due to these problems, we do not present age estimation at this point of our investigation.

#### 4.1.2 Correction for the Assumptions of IBD Length Distributions

The IBD length distribution was derived from sharing between two individuals, but we consider IBD sharing among many individuals and compute the raw IBD segment length as the maximal IBD sharing of any two individuals that possess the IBD segment. This results in overestimation of the lengths, because it is the maximum of all pairwise sharings. We also observed a second cause for raw IBD segments being longer than expected by the exponential distribution. The more individuals share an IBD segment, the more likely it is to find two individuals that share random minor alleles which would falsely extend the IBD segment.

Therefore, we corrected the raw lengths of IBD segments by locating the first tagSNV from the left (upstream) which is shared by at least 3/4 of the individuals that possess the IBD segment. This tagSNV is the left break point for the IBD segment. Analogously, we determined the right break point by the first tagSNV from the right (downstream) that is shared by at least 3/4 of the individuals. The distance between these break points is the (corrected) length of an IBD segment.

#### **4.1.3** Length Correction for IBD with Ancient Genomes

We are interested in IBD between modern human and ancient genomes. However, the human IBD segment length is not an appropriate measure for the length of IBD with ancient genomes because only a part of the IBD segment may match an ancient genome (see Figure 64).

We corrected the IBD segment lengths to obtain the length of the “ancient part” that matches a particular ancient genome. This “ancient part” must contain at least 8 tagSNVs, which is the minimum number of tagSNVs per IBD segments. First, the left (upstream) break point of the “ancient part” of an IBD segment genome was detected. This left break point is defined as the first location in the IBD segment from the left (upstream), where at least 4 out of 8 tagSNVs match the ancient genome. From the right (downstream), the right break point of the “ancient part” of an IBD segment was detected analogously. Since not all bases of the ancient genomes were called, we modified the definition of the break points and required at least 6 bases of the 8 tagSNVs to be called of which 3, have to match the ancient genome. If either the left or right break point of an “ancient part” could not be found, then this IBD segment does not contain an “ancient part” and was excluded from all further analyses.

Matching of an IBD segment and an ancient genome for IBD segment lengths analyses was defined as:

1. at least 15% of the tagSNVs of the IBD segment must match the ancient genome,
2. the “ancient part” of the IBD segment must contain at least 8 tagSNVs, and
3. 30% of the tagSNVs in the “ancient part” of the IBD segment must match the ancient genome.

### 4.2 Histograms of Lengths of IBD Segments for the Different Genomes

Figure 39 shows the histograms of IBD segment lengths for all IBD segments on the autosomes (human genome) and for IBD segments that match the Neandertal genome. For the human genome, a peak at 24,000 bp is visible, whereas, for the Neandertal genome, peaks are at 6,000 and 22,000 bp. It can be seen that IBD segments that match the Neandertal genome are shorter, thus also older.

**Figure 39:**
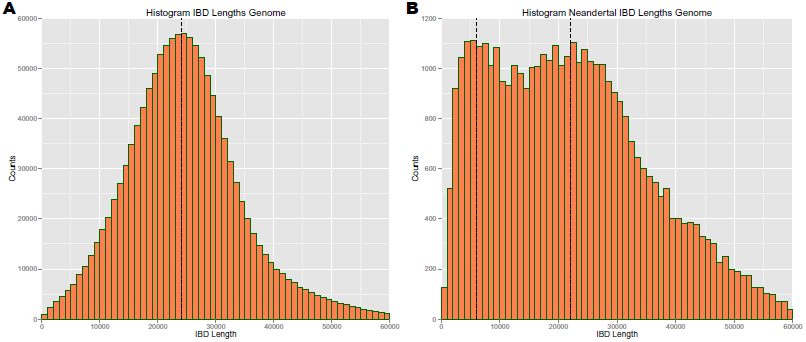
**Panel A**: Histogram of the IBD segment lengths for all IBD segments found in the 1000 Genomes Project data (human genome). The global peak is at 24,000 bp. **Panel B**: Histogram of the IBD segment lengths for IBD segments that match the Neandertal genome. Peaks at 6,000 bp and 22,000 bp are indicated.

Figure 40 shows the histograms of IBD segment lengths for all IBD segments that match the Denisova and the “Archaic” genome (“Archaic genome” contains IBD segments that match both the Denisova and Neandertal genome). For the Denisova genome, we have a peak at 5,000 bp, whereas, for the Archaic genome the peak is at 6,000 bp.

**Figure 40:**
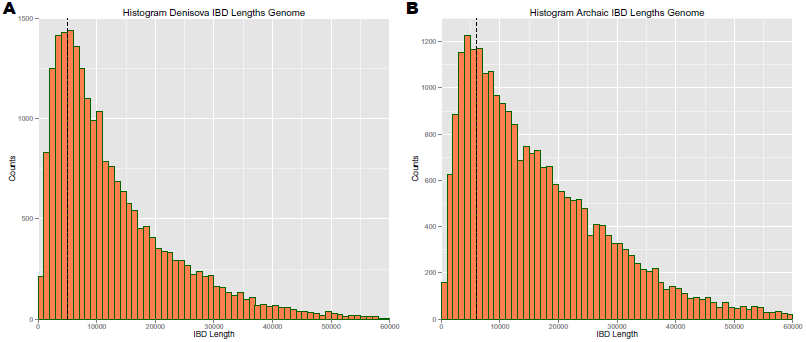
**Panel A**: Histogram of the IBD segment lengths for IBD segments that match the Denisova genome. A peak at 5,000 bp is indicated. **Panel B**: Histogram of the IBD segment lengths for IBD segments that match both the Neandertal and the Denisova genome (“Archaic genome”). The peak at 6,000 bp is indicated.

For chromosome X, Figure 41 shows the histograms of IBD segment lengths for all IBD segments (human genome) and for IBD segments that match the Neandertal genome. For the human genome, a peak can be seen at 33,000 bp, whereas, for the Neandertal genome, peaks are at 7,000, 15,000 and 39,000 bp. In general IBD segments that match the Neandertal genome are shorter, thus also older.

**Figure 41:**
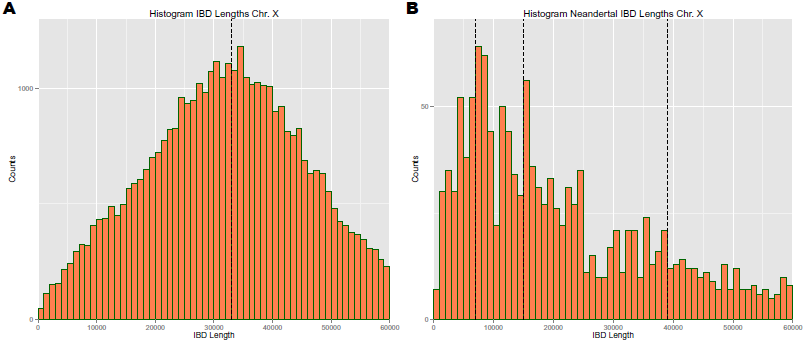
**Panel A**: Histogram of the IBD segment lengths for all IBD segments found on chromosome X of the 1000 Genomes Project data (human genome). The global peak is at 33,000 bp. **Panel B**: Histogram of the IBD segment lengths for IBD segments on chromosome X that match the Neandertal genome. Peaks at 7,000 bp, 15,000, and 39,000 bp are indicated.

Figure 42 shows the histograms of IBD segment lengths for all IBD segments on chromosome X that match the Denisova and the “Archaic” genome (“Archaic genome” contains IBD segments that match both the Denisova and Neandertal genome). For the Denisova genome, we have a peak at 6,000 bp, whereas, for the Archaic genome the peaks are at 8,000 and 19,000 bp.

**Figure 42:**
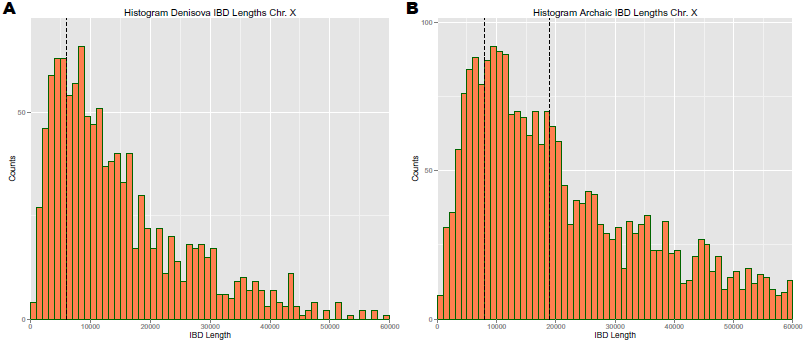
**Panel A**: Histogram of the IBD segment lengths for IBD segments that match the Denisova genome. The peak at 6,000 bp is indicated. **Panel B**: Histogram of the IBD segment lengths for IBD segments on chromosome X that match both the Neandertal and the Denisova genome (“Archaic genome”). Peaks at 8,000 bp and 19,000 bp are indicated.

Comparing the lengths of IBD segments on chromosome X with those on the autosomes, IBD segments on the X chromosome are on average longer. The recombination rate on chromosome X differs from the according rates of the autosomes. Per generation 2 out of 3 inherited X chromo-somes (the female ones) have a chance to recombine (29). This chance is given by the female recombination rate on chromosome X which is approximately equal to the autosomal recombination rate (26). Consequently, the sex-averaged recombination rate for chromosome X is 2/3 of the one for autosomes. Since the average IBD segment length depends inversely proportionally on the recombination rate, we can explain the differences of the global lengths peaks between chromosome X and the autosomes. On autosomes the global IBD segment length peak is at 24,000 bp, while on chromosome X it is at 33,000 bp, which roughly corresponds to the factor 2/3 introduced by the recombination rate difference. IBD segment lengths on the X chromosome confirm that our lengths estimations are inversely proportional to the number of recombinations, therefore in turn also to the number of generations or, equivalently, to the age of the segments.

IBD segment sharing between humans, Neandertals, and Denisovans may have two different reasons. First, the IBD segments may stem from a common ancestor and are passed on in each of these hominid groups. Secondly, IBD segment sharing may be caused by an introgression of one hominid group into another. For the former, the IBD segments are supposed to be on average shorter than for the latter scenario. Therefore, the peaks at shortest, and hence oldest, IBD segments must be assumed to stem from a common ancestor of Neandertals and Denisovans. Some of these IBD segments may have been lost either in Neandertals or Denisovans (at least in the specimens analyzed) and are therefore not attributed to the Archaic genome but to one of the other two groups. Introgression of Neandertals into the Denisova genome or vice versa and a subsequent gene flow into humans can explain long, therefore more recent, IBD segments that are attributed to the Archaic genome. This hypothesis is supported by the results of Prüfer et al. (39) which show evidence of Neandertal gene flow into Denisovans. Another explanation is that Neandertals and Denisovans did not differ at some DNA segments and one of them introduced the shared segments into the human genome.

### 4.3 IBD Segment Lengths of Human Populations

For all of the following analyses we excluded Admixed American individuals from the group of individuals that share an IBD segment in order to avoid influences on length distributions based on their admixed ancestry. Figure 43 shows the density of the lengths of IBD segments that are private to Asians vs. the density of the lengths of IBD segments that are private to Europeans. Since IBD segments are private to each continental population, the densities are based on disjoint IBD segment sets. Both show peaks at similar lengths. Asians show a global peak at 23,000 bp, while Europeans show a global peak at 24,000 bp. Both have smaller peaks between 40,000 bp and 50,000 bp.

**Figure 43:**
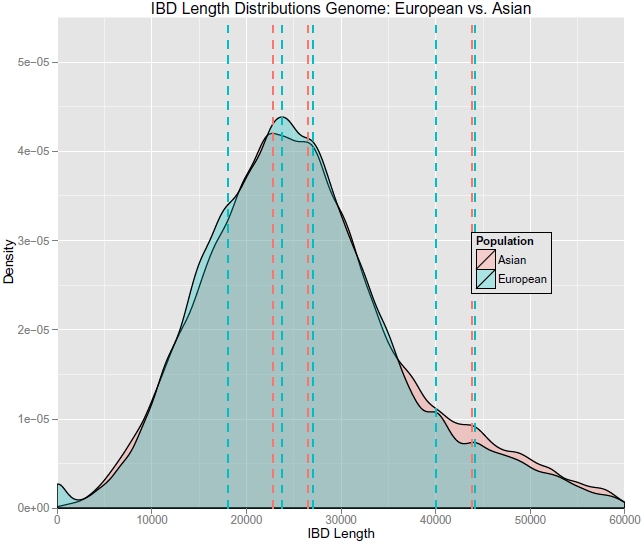
Density of lengths of IBD segments that are private to Asians vs. the analogous density for Europeans. Interesting peaks are marked by dashed lines. Asians have a global peak at 23,000 bp (red), while Europeans have the global peaks at 24,000 bp (blue). Both have smaller peaks around 45,000 bp.

Figure 44A shows the density of lengths of IBD segments that are private to Asians vs. lengths of IBD segments that are only shared between Asians and Africans. Figure 44B shows the same plot for Europeans instead of Asians. Hardly any difference is visible in the global peaks of length distributions of IBD segments that are private to a continental population (red dashed lines) and those shared with Africans (blue dashed lines): 23,000 vs. 26,000 bp for Asians and 24,000 vs. 23,000 bp for Europeans. However, segments that are also shared with Africans show an enrichment at shorter lengths compared to segments that are private to either Asians or Europeans. This enrichment is especially prominent for segments shared between Africans and Asians.

**Figure 44:**
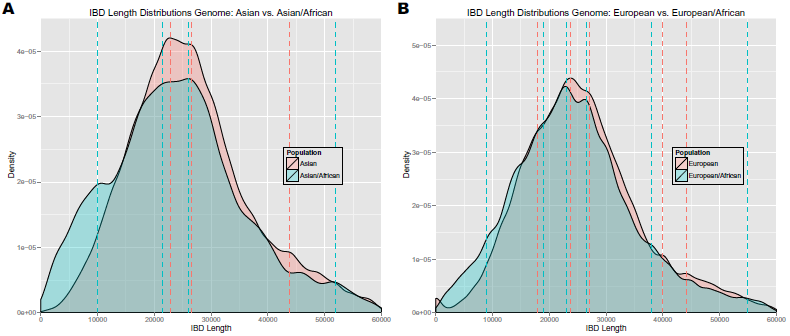
**Panel A**: Density of lengths of IBD segments that are private to Asians vs. density of IBD segment lengths shared only by Asians and Africans. **Panel B**: Density of lengths of IBD segments that are private to Europeans vs. density of IBD segment lengths shared only by Europeans and Africans. African-Asian IBD segments have peaks between 21,500 bp and 26,000 bp, as well as smaller peaks at 10,000 bp and 52,000 bp (blue dashed lines in panel B). African-European IBD segments have major peaks between 19,500 bp and 24,000 bp and smaller peaks at 9,000 bp, 38,000 bp, and 55,000 bp (blue dashed lines in panel B). The global Asian peak is at 23,000 bp (red dashed line in panel A), while the global peak for Europeans is at 24,000 bp (red dashed line in panel B). Both have smaller peaks around 45,000 bp.

Figure 45A shows the density of lengths of IBD segments that are private to Asians vs. lengths of IBD segments that are shared between Asians, Europeans, and Africans. Figure 45B shows the same plot for Europeans instead of Asians. Of course, the peaks for Asians and Europeans are the same as in Figure 44. Again, segments that are shared by all three continental populations show an enrichment at shorter lengths compared to segments that are private to either Asians or Europeans. We already assumed that IBD segments that are shared by all population groups predate the Out-of-Africa split and therefore have to be very old. These results confirm our assumptions since we show that these segments are very short.

**Figure 45:**
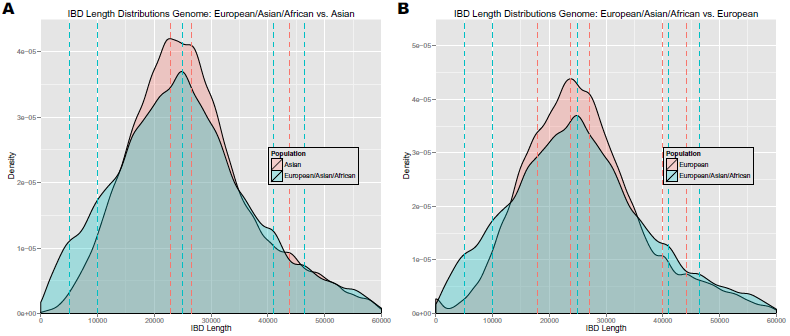
**Panel A**: Density of lengths of IBD segments that are private to Asians vs. density of IBD segment lengths shared by Asians, Europeans, and Africans. **Panel B**: Density of lengths of IBD segments that are private to Europeans vs. density of IBD segment lengths shared by Asians, Europeans, and Africans. Segments that are shared by all three continental populations show an enrichment at shorter lengths compared to segments that are private to either Asians or Europeans (blue vs. red).

Next, we investigated the effect on the length distribution if IBD segments are removed that are shared by Africans. Figure 46A shows the density of lengths of IBD segments that are private to Asians vs. the density of IBD segment lengths shared by Asians and Europeans, but not by Africans. In Figure 46B, the same plot as in Figure 46A is shown, but now compared to IBD segments that are private to Europeans. In Figure 45A and B, the density for IBD segments that are shared with Africans, has high values for shorter IBD segments (blue). In Figure 46A and B, this range of high values vanishes, because IBD segments that are shared with Africans are removed. A higher density region at longer segment length becomes visible for IBD segments that are shared by Europeans and Asians, but not by Africans compared to segments that are private to either Asians or Europeans.

**Figure 46:**
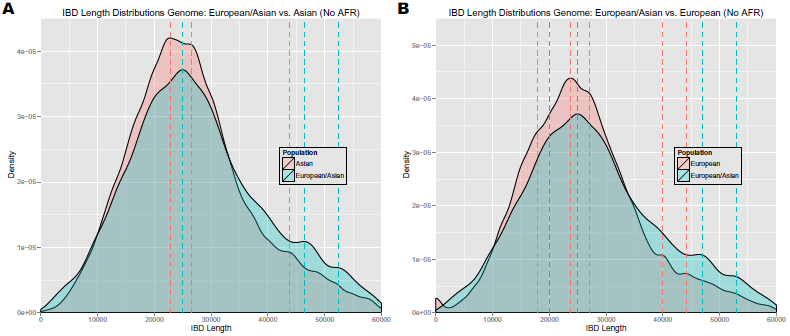
**Panel A**: Density of lengths of IBD segments that are private to Asians vs. density of IBD segment lengths shared by Asians and Europeans, but not by Africans meaning African-sharing IBD segments are removed as opposed to Figure 45A. **Panel B**: The same plot as in panel A, but IBD segments that are shared by Europeans and Asians are compared to IBD segments that are private to Europeans. African-sharing IBD segments are removed as opposed to Figure 45B. Both panels show that removing IBD segments shared with Africans in turn removes the higher densities at lower segment lengths that were present in Figure 45. Instead the densities for longer segment lengths are higher for IBD segments shared between Europeans and Asians (no Africans) compared to segments that are private to either Asians or Europeans.

Figure 47 shows the density of the lengths of IBD segments on chromosome X that are private to Asians vs. the density of the lengths of IBD segments that are private to Europeans. Since IBD segments are private to each continental population, the densities are based on disjoint IBD segment sets. Compared to the same densities for the autosomes, peaks are shifted towards longer segment lengths (see above for an explanation). Europeans have their global peak at 30,500 bp for chromosome X while it is at 24,000 bp for the autosomes. Asians have major peaks at 24,000 bp, 34,500 bp, and 50,500 bp for chromosome X compared to a single global peak at 23,000 bp for chromosomes 1-22.

**Figure 47:**
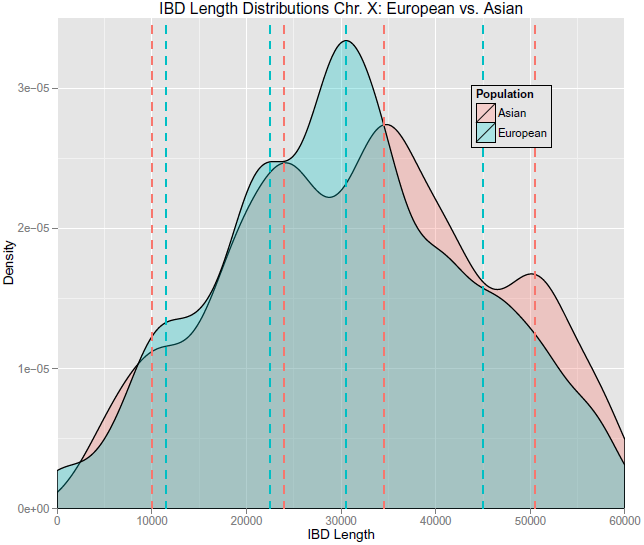
Density of lengths of IBD segments on chromosome X that are private to Asians vs. the analogous density for Europeans. Interesting peaks are marked by dashed lines. Europeans have their global peak at 30,500 bp (blue). Asians have major peaks at 24,000 bp, 34,500 bp, and 50,500 bp (red).

Figure 48A shows the density of lengths of IBD segments on chromosome X that are private to Asians vs. lengths of IBD segments that are only shared between Asians and Africans. Figure 48B shows the same plot for Europeans instead of Asians. The African-Asian density peaks are at similar lengths as the Asian peaks at 24,000 bp, 34,500 bp, and also at 46,000 bp. An additional peak at 10,000 bp is more prominent for segments shared by Africans and Asians. This high density region at shorter segment lengths is not as prominent as for the autosomes. African-European IBD segments have a major peak 45,000 bp and no enrichment of shorter segments compared to private European IBD segments.

**Figure 48:**
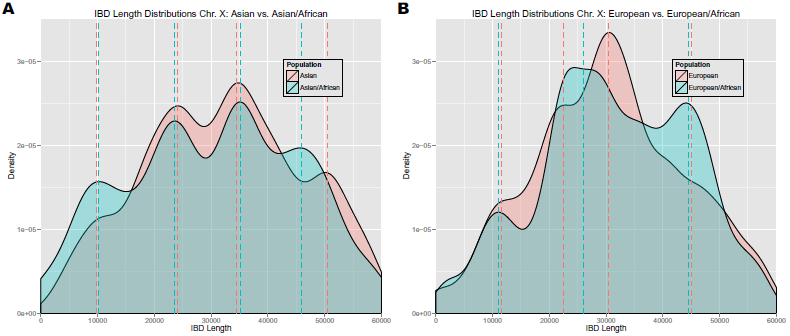
**Panel A**: Density of lengths of IBD segments on chromosome X that are private to Asians vs. density of IBD segment lengths shared only by Asians and Africans. **Panel B**: Density of lengths of IBD segments on chromosome X that are private to Europeans vs. density of IBD segment lengths shared only by Europeans and Africans. The African-Asian density peaks (blue dashed lines in panel A) are at similar lengths as the Asian peaks (red dashed lines in panel A) at 24,000 bp, 34,500 bp, and also at 46,000 bp. An additional peak at 10,000 bp is more prominent for segments shared by Africans and Asians. In contrast to that African-European IBD segments (blue dashed lines in panel B) have a major peak 45,000 bp and no enrichment of shorter segments compared to private European IBD segments (red dashed line in panel B).

Figure 49A shows the density of lengths of IBD segments on chromosome X that are private to Asians vs. lengths of IBD segments that are shared between Asians, Europeans, and Africans. Figure 49B shows the same plot for Europeans instead of Asians. Of course, the peaks for Asians and Europeans are the same as in Figure 48. As for the autosomes, segments that are shared by all three continental populations show an enrichment at shorter lengths compared to segments that are private to either Asians or Europeans. This enrichment is much weaker on chromosome X.

**Figure 49:**
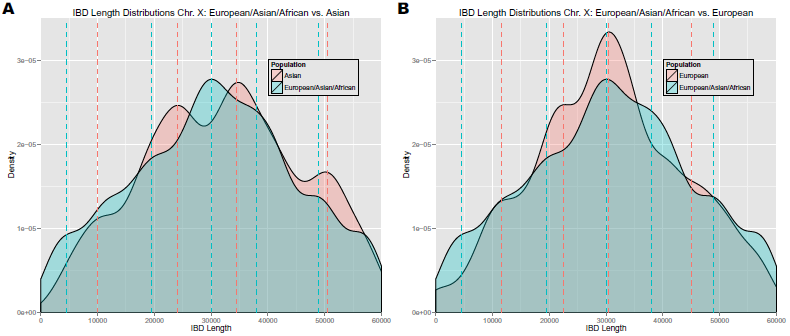
**Panel A**: Density of lengths of IBD segments on chromosome X that are private to Asians vs. density of IBD segment lengths shared by Asians, Europeans, and Africans. **Panel B**: Density of lengths of IBD segments on chromosome X that are private to Europeans vs. density of IBD segment lengths shared by Asians, Europeans, and Africans. Segments that are shared by all three continental populations show a slight enrichment at shorter lengths compared to segments that are private to either Asians or Europeans (blue vs. red). The enrichment is not as prominent as for the autosomes.

Next, we investigated the effect on the length distribution if IBD segments are removed that are shared by Africans. Figure 50A shows the density of lengths of IBD segments on chromosome X that are private to Asians vs. the density of IBD segment lengths shared by Asians and Europeans, but not by Africans. In Figure 50B, the same plot as in Figure 50A is shown, but now compared to IBD segments that are private to Europeans. In Figure 49A and B, the density for IBD segments that are shared with Africans, has high values for shorter IBD segments (blue). In Figure 50A and B, this range of high values vanishes, because IBD segments that are shared with Africans are removed.

**Figure 50:**
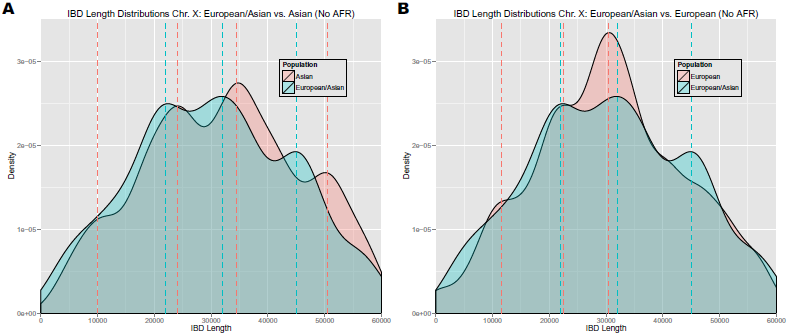
**Panel A**: Density of lengths of IBD segments on chromosome X that are private to Asians vs. density of IBD segment lengths shared by Asians and Europeans, but not by Africans meaning African-sharing IBD segments are removed as opposed to Figure 49A. **Panel B**: The same plot as in panel A, but IBD segments that are shared by Europeans and Asians are compared to IBD segments that are private to Europeans. African-sharing IBD segments are removed as opposed to Figure 49B. Both panels show that removing IBD segments shared with Africans in turn reduces the higher densities at lower segment lengths that were present in Figure 49.

### 4.4 Lengths of IBD Segments that Match the Denisova or Neandertal Genome

#### 4.4.1 Lengths of IBD Segments that Match the Denisova Genome

Figure 51A shows the density of the lengths of all IBD segments on chromosomes 1-22 (human genome) vs. the density of the lengths of IBD segments that match the Denisova genome. The length density of all IBD segments has a peak at 24,000 bp, while the length distribution of IBD with the Denisova genome has a peak at 4,500 bp. Clearly, segments of IBD with the Denisova genome are shorter and therefore older than those solely shared among present day humans (red density above blue at the left hand side). We were interested in how different continental populations share the Denisova genome. Figure 51B shows densities of lengths of IBD segments that match the Denisova genome and are enriched by a particular continental population. For all populations combined as well as for Africans and Europeans, the global density peak is visible between 4,500 and 5,500 bp. Europeans have additional peaks at 14,000 bp and 22,500 bp, as well as between 30,000 and 45,000 bp. Denisova-matching segments that are shared by Asians are generally longer with a global peak at 24,000 bp and additional smaller peaks at 13,000 bp, 18,000 bp, 40,000 bp, and 50,500 bp. These densities seem to reveal that there was a gene flow from Denisovans into the Asian genome outside Africa. However, also Europeans show some hints of introgression from the Denisovans after migration out of Africa.

**Figure 51:**
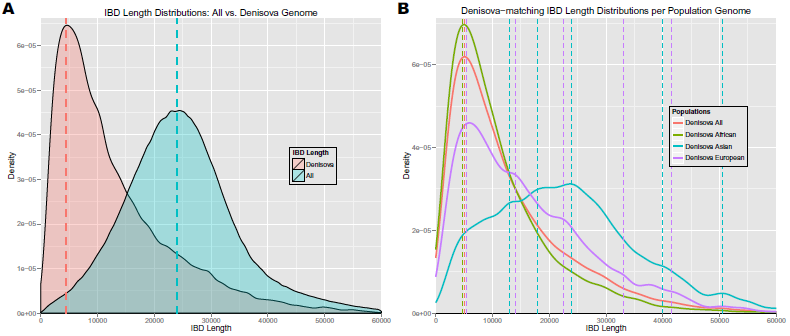
**Panel A**: Density of the lengths of all IBD segments vs. the lengths of IBD segments that match the Denisova genome. The dashed lines indicate the peaks at 9,000 bp for the Denisova and 24,000 bp for the human genome. **Panel B**: Densities of lengths of IBD segments that match the Denisova genome and are enriched in a particular population. The dashed lines indicate density peaks. For all populations combined as well as for Africans and Europeans, the global density peak is visible between 4,500 and 5,500 bp. Europeans have additional peaks at 14,000 bp and 22,500 bp, as well as between 30,000 and 45,000 bp. Denisova-matching segments that are shared by Asians are generally longer with a global peak at 24,000 bp and additional smaller peaks at 13,000 bp, 18,000 bp, 40,000 bp, and 50,500 bp.

These peaks can be seen more clearly if only IBD segments are considered that are private to a population. Figure 52A shows densities of lengths of IBD segments that match the Denisova genome and are private to a population. For all populations together, and for Africans alone, the peaks of the length densities are around 5,500 bp. For Denisova-matching IBD segments private to Europeans the peak is at 13,700 bp and for IBD segments private to Asians, it is at 23,200 bp. The density of IBD segment lengths that match the Denisova genome and are private to Asians is also high around 40,000 bp and 50,000 bp. Europeans have an additional peak at 25,000 bp and 38,800 bp. Figure 52B shows densities of lengths of IBD segments that match the Denisova genome and are private to Africans vs. IBD segments that are not observed in Africans. The global peak for Africans is at 4,500 bp, while the density of lengths of IBD segments that are not observed in Africans has a global peak at 22,000 bp. Africans have shorter and therefore older segments probably stemming from common ancestors of Denisovans and humans. For non-African populations, the high densities for longer IBD segments hint at an introgression from Denisovans after migration out of Africa.

**Figure 52:**
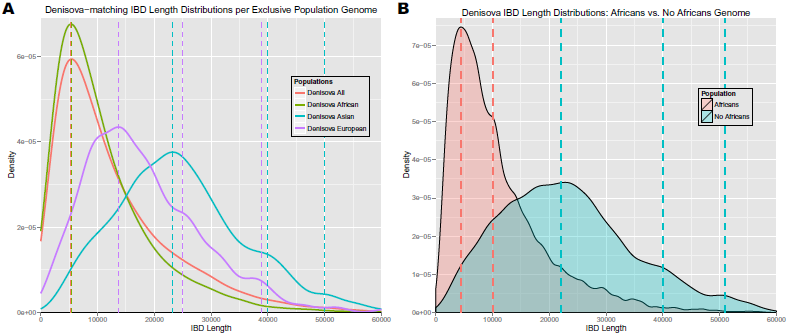
**Panel A**: Densities of lengths of IBD segments that match the Denisova genome and are private to a population. The peaks are around 5,500 bp for all IBD segments matching the Denisova genome and for Africans alone, 13,700 for Europeans, and 23,200 for Asians. **Panel B**: Densities of lengths of IBD segments that match the Denisova genome and are private to Africans vs. IBD segments that are not observed in Africans. The global peak for Africans is at 5,500 bp, while the density of lengths of IBD segments that are not observed in Africans has a global peak at 22,000 bp.

Figure 53A shows the density of the lengths of all IBD segments on chromosome X (human genome) vs. the density of the lengths of IBD segments that match the Denisova genome. The length density of all IBD segments has a peak at 34,000 bp, while the length distribution of IBD with the Denisova genome has a peak at 6,000 bp. Clearly, segments of IBD with the Denisova genome are shorter and therefore older than those solely shared among present day humans (red density above blue at the left hand side). We were interested in how different populations share the Denisova genome. Figure 53B shows densities of lengths of IBD segments that match the Denisova genome and are enriched by a particular continental population. The density for all populations combined matches the density for Africans because almost all Denisova-matching segments are shared by at least one African individual. They have a global density peak around 6,200 bp and an additional peak at 28,000 bp. Europeans and Asians have global peaks at 5,300 bp and 8,400 bp, respectively, as well as additional smaller peaks between 15,000 and 35,000 bp. The European and Asian density peaks are based on very few IBD segments and therefore should be considered with caution. No clear signal of introgression can be observed on the X chromosome as a strong peak at longer segment length is missing.

**Figure 53:**
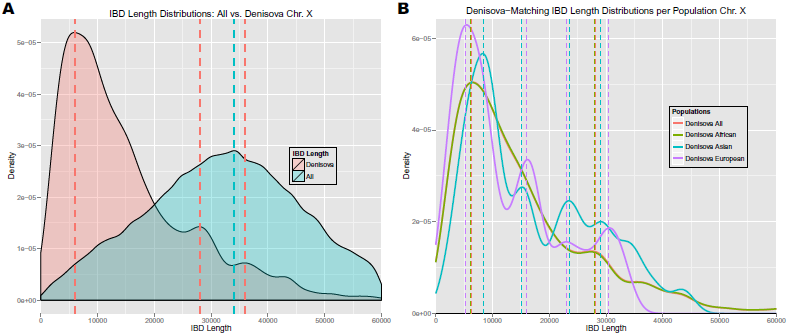
**Panel A**: Density of the lengths of all IBD segments on chromosome X vs. the lengths of IBD segments that match the Denisova genome. The dashed lines indicate the peaks at 6,000 bp for the Denisova and 34,000 bp for the human genome. **Panel B**: Densities of lengths of IBD segments on chromosome X that match the Denisova genome and are enriched in a particular population. The dashed lines indicate density peaks. The density for all populations combined matches the density for Africans because almost all Denisova-matching segments are shared by at least one African individual. They have a global density peak around 6,200 bp and an additional peak at 28,000 bp. Europeans and Asians have global peaks at 5,300 bp and 8,400 bp, respectively, as well as additional smaller peaks between 15,000 and 35,000 bp.

Figure 54A shows densities of lengths of IBD segments on chromosome X that match the Denisova genome and are private to a population. For all populations together, and for Africans alone, the peak of the length density is around 7,000 bp with Africans having an additional peak at 28,000 bp. For IBD segments private to Europeans, the global peak is at 4,000 bp and an additional peak at 10,000 bp. IBD segments private to Asians have density peaks at 5,500 bp, 16,000 bp, 24,000 bp, and 35,000 bp. Here the number of IBD segments private to either Europeans or Asians is very small (below 20). Therefore, the peaks are probably at random length positions and analyses based on them should not be trusted. Figure 54B shows densities of lengths of IBD segments that match the Denisova genome and are private to Africans vs. IBD segments that are not observed in Africans. The global peak for Africans is at 6,200 bp and an additional peak is visible at 28,000 bp, while the density of lengths of IBD segments that are not observed in Africans has a global peak at 5,700 bp and an additional peak at 32,000 bp. These results should be considered with care, because the number of non-African IBD segments is low. In contrast to the autosomes, there is no clear abundance of longer Denisova-matching IBD segments on the X chromosome that are shared by non-African individuals.

**Figure 54:**
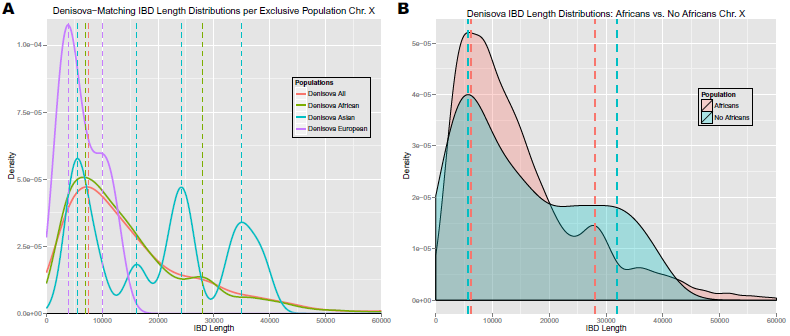
**Panel A**: Densities of lengths of IBD segments on chromosome X that match the Denisova genome and are private to a population. For all populations together, and for Africans alone, the peaks of the length densities are around 7,000 bp with Africans having an additional peak at 28,000 bp. For IBD segments private to Europeans the global peak is at 4,000 bp and an additional peak is at 10,000 bp. IBD segments private to Asians have density peaks are at 5,500 bp, 16,000 bp, 24,000 bp, and 35,000 bp. The number of IBD segments private to either Europeans or Asians is very small, therefore these peaks should not be trusted. **Panel B**: Densities of lengths of IBD segments on chromosome X that match the Denisova genome and are private to Africans vs. IBD segments that are not observed in Africans. The global peak for Africans is at 6,200 bp and an additional peak is visible at 28,000 bp, while the density of lengths of IBD segments that are not observed in Africans has a global peak at 5,700 bp and an additional peak at 32,000 bp.

#### 4.4.2 Lengths of IBD Segments that Match the Neandertal Genome

Figure 55A shows the density of the lengths of all IBD segments on the autosomes (human genome) vs. the density of the lengths of IBD segments that match the Neandertal genome. The length density of all IBD segments has a peak at 24,000 bp, while the length distribution of IBD segments with the Neandertal genome (the part of the IBD segment that matches the Neandertal genome) has peaks at 5,500 bp, 23,000 bp, and 43,000 bp. Figure 55B shows densities of lengths of IBD segments that match the Neandertal genome and are enriched in a particular continental population. The peaks of the length distribution for IBD segments that are shared by all populations combined are at 5,500 bp and 24,000 bp. Africans have their global peak also around 5,000 bp. Europeans and Asians have their global peaks around 25,000 bp and additional peaks around 40,000 bp. The density peak for Africans is clearly separated from the density peaks for Europeans and Asians which almost match each other. This hints at introgression from the Neandertals into anatomically modern humans that were the ancestors of Europeans and Asians after these humans left Africa. The higher density of short IBD segments which are prominent in Africans in the range of 5,000–20,000 bp hints at old DNA segments that humans share with the Neandertal genome. Again, our results contradict the hypothesis of Prüfer et al. (39) that Neandertal ancestry in Africans is due to back-to-Africa admixture, but instead hint at an earlier origin from within Africa. These short and old DNA segments may even stem from common ancestors of humans and Neandertals.

**Figure 55:**
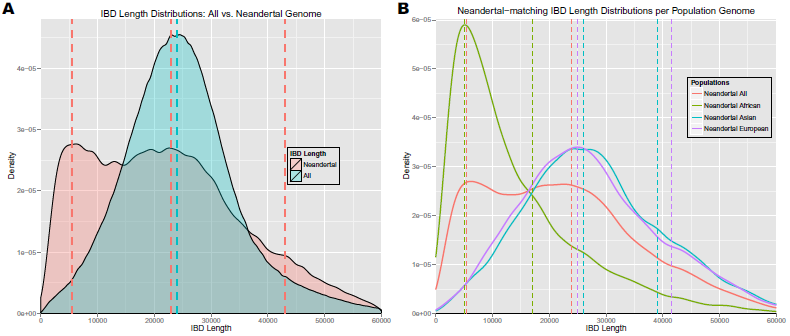
**Panel A**: Density of the lengths of all IBD segments vs. the lengths of IBD segments that match the Neandertal genome. The dashed lines indicate the peaks of densities at 24,000 bp for all IBD segments (blue) and at 5,500 bp, 23,000 bp, and 43,000 bp for Neandertal-matching segments (red). **Panel B**: Densities of lengths of IBD segments that match the Neandertal genome and are enriched in a particular population. The dashed lines indicate the density peaks around 5,000 bp and 17,000 bp for Africans, 5,500 bp and 24,000 bp for all populations combined, 25,000 bp and 41,500 bp for Europeans, and 26,000 bp, and 39,000 bp for Asians.

Next, we considered IBD segments that are private to a continental population. Figure 56A shows densities of lengths of IBD segments that match the Neandertal genome and are private to a population. The peaks are at 5,500 bp for Africans and at 6,000 bp and 24,000 bp for all humans combined. The density for Europeans has peaks at 23,500 bp, and 43,500 bp. Asians have density peaks at 23,000 bp, and 28,500 bp. The densities of Asians and Europeans agree well with each other. They have peaks at longer segment lengths compared to Africans. Therefore, we were interested in IBD segments that match the Neandertal genome and that are private to Africans and those which are not observed in Africans hence shared by either Asians or Europeans or both. Figure 56B shows densities of lengths of IBD segments that match the Neandertal genome and are private to Africans vs. IBD segments that are not observed in Africans. The peaks for Africans are around 5,000 bp and 16,000 bp, while IBD segments that are not observed in Africans have a global length density peak at 24,000 bp and a smaller peak at 43,000 bp. Most prominently, non-African IBD segments that match the Neandertal genome are enriched in regions of longer segment lengths.

**Figure 56:**
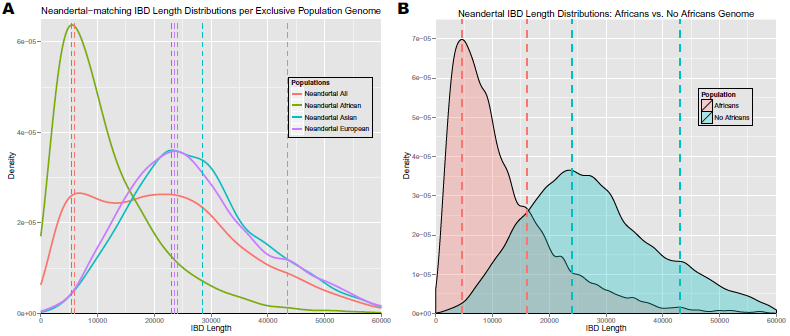
**Panel A**: Densities of lengths of IBD segments that match the Neandertal genome and are private to a population. The major peaks are at 5,500 bp for Africans, 6,000 bp and 24,000 bp for all humans combined, 23,500 bp for Europeans, and 23,000 bp for Asians. **Panel B**: Densities of lengths of IBD segments that match the Neandertal genome and are private to Africans vs. IBD segments that are not observed in Africans. The peaks for Africans are around 5,000 bp and 16,000 bp, while IBD segments that are not observed in Africans have a global length density peak at 24,000 bp and a smaller peak at 43,000 bp. Most prominently, non-African IBD segments that match the Neandertal genome are enriched in regions of longer segment lengths (blue density).

Figure 57A shows the density of the lengths of all IBD segments on chromosome X (human genome) vs. the density of the lengths of IBD segments that match the Neandertal genome. The length density of all IBD segments has a peak at 34,000 bp, while the length distribution of IBD segments with the Neandertal genome (the part of the IBD segment that matches the Neandertal genome) has peaks at 7,500 bp, 15,000 bp, 23,000 bp, and 35,000 bp. Figure 57B shows densities of lengths of IBD segments that match the Neandertal genome and are enriched in a particular continental population. The peaks of the length distribution for IBD segments that are shared by all populations combined are at 7,500 bp, 15,000 bp, and 35,000 bp. Africans have their global peak also around 7,500 bp and additional peaks at 15,000 bp and 33,000 bp. Europeans and Asians have peaks around 17,000 bp, 37,000 bp, and 51,000 bp. The density peak for Africans is clearly separated from the global density peaks for Europeans and Asians which almost match each other. Therefore, also on chromosome X there are hints at introgression from the Neandertals into anatomically modern humans that were the ancestors of Europeans and Asians after these humans left Africa. Very old and short DNA segments that stem from ancestors of humans and Neandertals have survived on chromosome X of African individuals.

**Figure 57:**
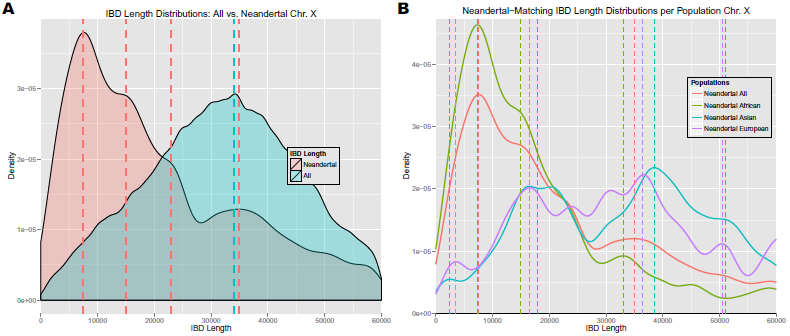
**Panel A**: Density of the lengths of all IBD segments on chromosome X vs. the lengths of IBD segments that match the Neandertal genome. The dashed lines indicate the peaks of densities at 34,000 bp for all IBD segments (blue) and at 7,500 bp, 15,000 bp, 23,000 bp, and 35,000 bp for Neandertal-matching segments (red). **Panel B**: Densities of lengths of IBD segments on chro-mosome X that match the Neandertal genome and are enriched in a particular population. The dashed lines indicate the density peaks around 7,500 bp, 15,000 bp, and 35,000 bp for Africans and for all populations combined, and around 17,000 bp, 37,000 bp, and 51,000 bp for Europeans and Asians.

Next, we considered IBD segments on chromosome X that are private to a continental poputation. Figure 58A shows densities of lengths of IBD segments that match the Neandertal genome and are private to a population. The global peaks for Africans and all populations combined are around 8,000 bp. The length density for Europeans has major peaks at 19,000 bp, 40,000 bp, and 59,500 bp. Asians have major density peaks at 22,500 bp, 31,000 bp, 39,500 bp, and 48,500 bp. Again, the densities of Asians and Europeans are similar to each other. They have peaks at longer segment lengths compared to Africans. Therefore, we were interested in IBD segments that match the Neandertal genome and that are private to Africans opposed to those which are not observed in Africans hence shared by either Asians or Europeans or both. Figure 58B shows densities of lengths of IBD segments that match the Neandertal genome and are private to Africans vs. IBD segments that are not observed in Africans. The major peaks for Africans are around 7,500 bp, 15,000 bp and 33,000 bp, while IBD segments that are not observed in Africans have a global length density peak at 40,000 bp and a smaller peak at 21,000 bp. Most prominently, non-African IBD segments that match the Neandertal genome are enriched in regions of longer segment lengths.

**Figure 58:**
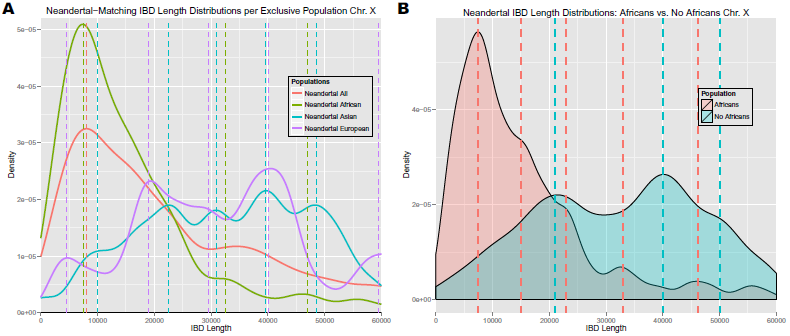
**Panel A**: Densities of lengths of IBD segments on chromosome X that match the Neandertal genome and are private to a population. For Africans and all populations combined the global peaks are around 8,000 bp. The density for Europeans has major peaks at 19,000 bp, 40,000 bp, and 59,500 bp. Asians have major density peaks at 22,500 bp, 31,000 bp, 39,500 bp, and 48,500 bp. **Panel B**: Densities of lengths of IBD segments on chromosome X that match the Neandertal genome and are private to Africans vs. IBD segments that are not observed in Africans. The major peaks for Africans are around 7,500 bp, 15,000 bp and 33,000 bp, while IBD segments that are not observed in Africans have a global length density peak at 40,000 bp and a smaller peak at 21,000 bp. Most prominently, non-African IBD segments that match the Neandertal genome are enriched in regions of longer segment lengths (blue density).

#### 4.4.3 Lengths of IBD Segments that Match Neandertal & Denisova

IBD segments that match the “Archaic genome” are IBD segments that match both the Neandertal and the Denisova genome. Segments matching the Archaic genome stem either from a genome of archaic hominids which were ancestors of Neandertals and Denisovans or they stem from introgression of one hominid group into another. Garrigan et al. (13) were the first to present evidence for a prolonged period of ancestral population subdivision followed by relatively recent interbreeding in the history of human populations. Later, Wall et al. (59) found evidence that suggests that admixture between diverged hominid groups may have been a common feature of human evolution. Recently Prüfer et al. (39) published findings that suggest gene flow from Neandertals into Denisovans as well as an additional ancestral component in Denisovans from an unknown ancient population.

Figure 59A shows the density of the lengths of all IBD segments (human genome) vs. the density of the lengths of IBD segments that match the Archaic genome. For the human genome, the global density peak is again at 24,000 bp. For IBD with the Archaic genome, the global density peak is at 5,000 bp, but smaller peaks at 15,500 bp and 23,500 bp can be observed. Figure 59B shows densities of lengths of IBD segments that match the Archaic genome and are enriched in a particular continental population. The global density peaks for all populations as well as for Africans can be seen around 5,000 bp. Europeans and Asians have peaks around 12,000 bp and 25,000 bp. Similar to IBD segments matching the Denisova or Neandertal genome separately, segments shared by Europeans or Asians are longer than segments shared by Africans.

**Figure 59:**
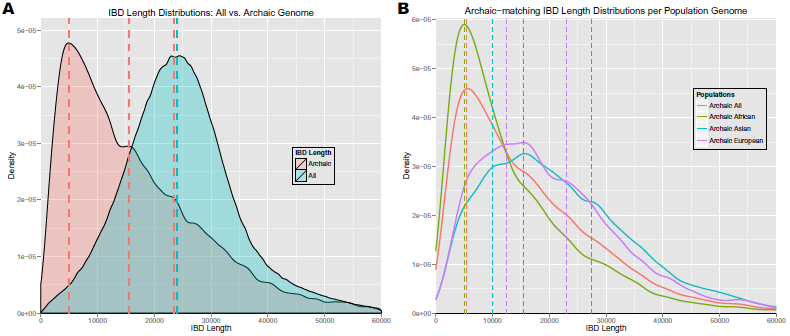
**Panel A**: Density of the lengths of all IBD segments vs. the lengths of IBD segments that match the Archaic genome. The dashed lines indicate the global peaks of densities at 5,000 bp and 25,000 bp as well as the smaller peaks at 15,500 bp and 23,500 bp. **Panel B**: Densities of lengths of IBD segments that match the Archaic genome and are enriched in a particular population. The dashed lines indicate the density peaks with the global peaks for all populations combined as well as for Africans around 5,000 bp. Europeans and Asians have peaks at longer segment lengths ranging from 10,000 bp to 30,000 bp.

Next, we considered IBD segments that are private to a continental population. Figure 60A shows densities of lengths of IBD segments that match the Archaic genome and are private to a population. All populations combined, as well as Africans alone have a peak around 6,000 bp. Asians and Europeans have their global peak between 10,000 bp and 18,000 bp. Figure 60B shows densities of lengths of IBD segments that match the Archaic genome and are private to Africans vs. IBD segments that are not observed in Africans. The global peak for Africans is at 4,500 bp and a smaller one at 15,000 bp, while lengths of IBD segments that are not observed in Africans have peaks at 10,000 bp and 15,500 bp. Most prominently, non-African IBD segments that match the Archaic genome are enriched at larger segment lengths. This enrichment seems to be caused by events after humans migrated out of Africa. The introgression from the Neandertal into ancestors of modern humans may also have introduced a part of the Denisova genome that has been contained in the Neandertal genome, or vice versa. We would consider this part of the human genome as matching the Archaic genome.

**Figure 60:**
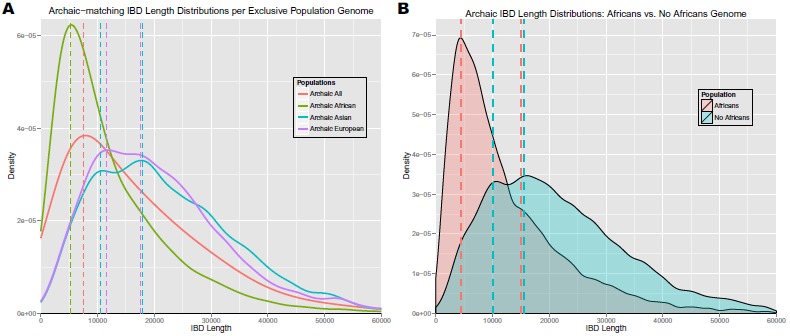
**Panel A**: Densities of lengths of IBD segments that match the Archaic genome and are private to a population. The peaks are around 7,000 bp and 10,000 bp for all populations combined, as well as for Africans alone. Asians and Europeans have their global peak between 10,000 bp and 18,000 bp. **Panel B**: Densities of lengths of IBD segments that match the Archaic genome and are private to Africans vs. IBD segments that are not observed in Africans. The global peak for Africans is at 4,500 bp and a smaller one at 15,000 bp, while lengths of IBD segments that are not observed in Africans have peaks at 10,000 bp and 15,500 bp. Non-African IBD segments that match the Archaic genome are enriched at larger segment lengths (blue density).

Figure 61A shows the density of the lengths of all IBD segments on chromosome X (human genome) vs. the density of the lengths of IBD segments that match the Archaic genome. For the human genome, the global density peak is again at 34,000 bp. For IBD with the Archaic genome, the global density peak is at 9,500 bp, but smaller peaks at 18,000 bp and 34,500 bp can be observed. Figure 61B shows densities of lengths of IBD segments that match the Archaic genome and are enriched in a particular continental population. All populations combined as well as each population separately reveal peaks around 9,000 bp, 19,000 bp, and 35,000 bp. Europeans and Asians have additional smaller peaks at longer segment lengths. On chromosome X, Archaic-matching segments shared by Europeans or Asians are not clearly longer than segments shared by Africans.

**Figure 61:**
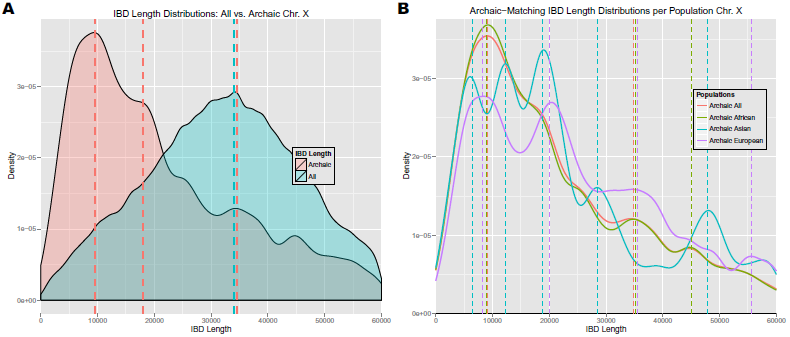
**Panel A**: Density of the lengths of all IBD segments on chromosome X vs. the lengths of IBD segments that match the Archaic genome. The dashed lines indicate the global peaks of densities at 34,000 bp and 9,500 bp as well as the smaller peaks at 18,000 bp and 34,500 bp. **Panel B**: Densities of lengths of IBD segments on chromosome X that match the Archaic genome and are enriched in a particular population. All populations combined as well as each population separately reveal peaks around 9,000 bp, 19,000 bp, and 35,000 bp. Europeans and Asians have additional smaller peaks at longer segment lengths.

Next, we considered IBD segments that are private to a continental population. Figure 62A shows densities of lengths of IBD segments on chromosome X that match the Archaic genome and are private to a population. All populations combined, as well as Africans alone have a global peak around 9,000 bp. Asians have larger peaks at 5,500 bp and 19,000 bp, as well as smaller peaks at longer segment lengths. Europeans have their larger peaks at 21,000 bp and between 32,000 bp and 37,000 bp. Again, these results have to be considered with care as only few Archaic-matching IBD segments on chromosome X are exclusively shared by a non-African population. Figure 62B shows densities of lengths of IBD segments that match the Archaic genome and are private to Africans vs. IBD segments that are not observed in Africans. The global peak for Africans is at 8,800 bp and smaller ones at 25,000 bp, 35,500 bp, and 45,000 bp, while lengths of IBD segments that are not observed in Africans have peaks at 7,500 bp, 18,500 bp, 30,000 bp, and 49,000 bp. Non-African IBD segments that match the Archaic genome seem to be enriched at larger segment lengths, although not as clearly as on the autosomes.

**Figure 62:**
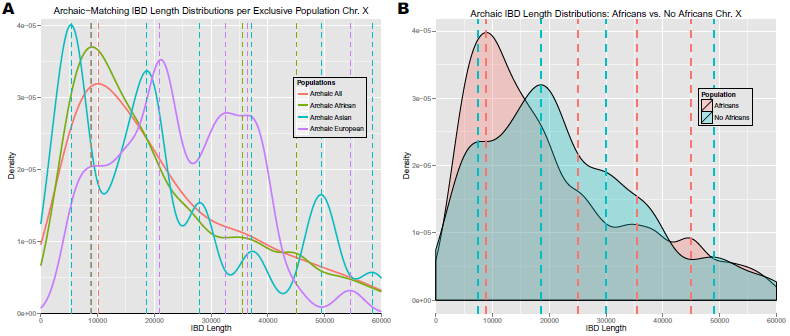
**Panel A**: Densities of lengths of IBD segments on chromosome X that match the Archaic genome and are private to a population. All populations combined, as well as Africans alone have global peaks around 9,000 bp. Asians have larger peaks at 5,500 bp and 19,000 bp, as well as smaller peaks at longer segment lengths. Europeans have their larger peaks at 21,000 bp and between 32,000 bp and 37,000 bp. Again, these results have to be considered with care as only few Archaic-matching IBD segments on chromosome X are exclusively shared by a non-African population. **Panel B**: Densities of lengths of IBD segments on chromosome X that match the Archaic genome and are private to Africans vs. IBD segments that are not observed in Africans. The global peak for Africans is at 8,800 bp and smaller ones at 25,000 bp, 35,500 bp, and 45,000 bp, while lengths of IBD segments that are not observed in Africans have peaks at 7,500 bp, 18,500 bp, 30,000 bp, and 49,000 bp. Non-African IBD segments that match the Archaic genome are enriched at larger segment lengths (blue density).

## 5 Revisiting the X Chromosome

The X chromosome is unique among the chromosomes because of its particular role in population genetics. It can be utilized to check the assumptions concerning the lengths distributions as its sex-averaged recombination rate differs from the autosomes. Per generation 2 out of 3 inherited X chromosomes (the female ones) have a chance to recombine (29). This chance is given by the female recombination rate on chromosome X which is approximately equal to the autosomal recombination rate (26). Consequently, the sex-averaged recombination rate for chromosome X is 2/3 of the one for autosomes. IBD segment lengths on the X chromosome confirm that our lengths estimations are inversely proportional to the number of recombinations (see Section 4.2). Thus, the number of generations or, equivalently, the age of the segments is inversely proportional to IBD segment lengths.

Furthermore, the IBD segments on the X chromosome shed additional light onto the demographic history of humans and ancient genomes. Since the recombination rate on chromosome X is 2/3 of that of the autosomes, we expect longer segments of the same age. For chromosome X, very old segments tend to be broken into small pieces to a lesser extent than for autosomes, therefore those old IBD segments are easier to detect. This is one explanation for the larger number of IBD segments of the X chromosome that match the Archaic genome and are shared by Africans (see Section 3.2.3). These segments must be assumed to stem from times where ancestors of these hominid groups either admixed in Africa or had a common ancestor, therefore they are indeed very old. The corresponding segments on autosomes are more difficult to find because they are much shorter.

We already mentioned that the X chromosome contains almost no long IBD segments that are shared between humans and Denisovans (see Section 4.4.1). This can be explained either by the fact that Denisova-DNA never entered the human X chromosome outside of Africa or on this chromosome the Denisova sequence got lost in humans. The former explanation would indicate that almost exclusively male Denisovans fathered male offspring with human females. The latter explanation would hint at some selective pressure. Along the same lines, the X chromosome contains fewer long IBD segments that are shared between the humans and Neandertals than the autosomes (see Section 4.4.2). However, still either more Neandertal DNA entered the X chromosome or survived therein compared to the Denisova DNA.

## 6 Examples of IBD Segments that Match Ancient Genomes

Figure 63 shows a typical example of an IBD segment that matches the Denisova genome and is shared exclusively among Asians. Figure 64 shows an IBD segment that matches the Denisova genome and is shared by Africans and one Admixed American. Figure 65 shows an example of an IBD segment that matches the Denisova genome and is shared by Europeans and Asians.

**Figure 63:**
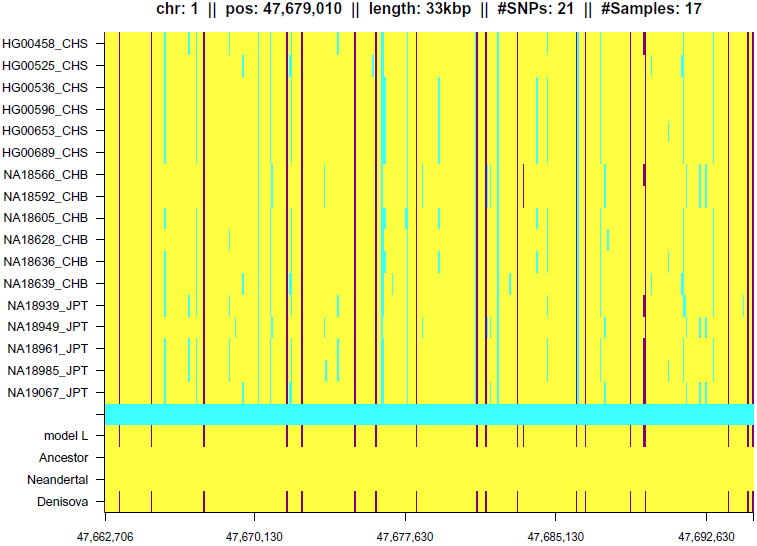
Example of an IBD segment matching the Denisova genome shared exclusively among Asians. The data analyzed by HapFABIA were genotypes of the 1000 Genomes Project. The rows give all individuals that contain the IBD segment and columns consecutive SNVs. Major alleles are shown in yellow, minor alleles of tagSNVs in violet, and minor alleles of other SNVs in cyan. The row labeled “model L” indicates tagSNVs identified by HapFABIA in violet. The rows “Ancestor”, “Neandertal”, and “Denisova” show bases of the respective genomes in violet if they match the minor allele of the tagSNVs (in yellow otherwise). For the “Ancestor genome”, we used the reconstructed common ancestor sequence that was provided as part of the 1000 Genomes Project data.

**Figure 64:**
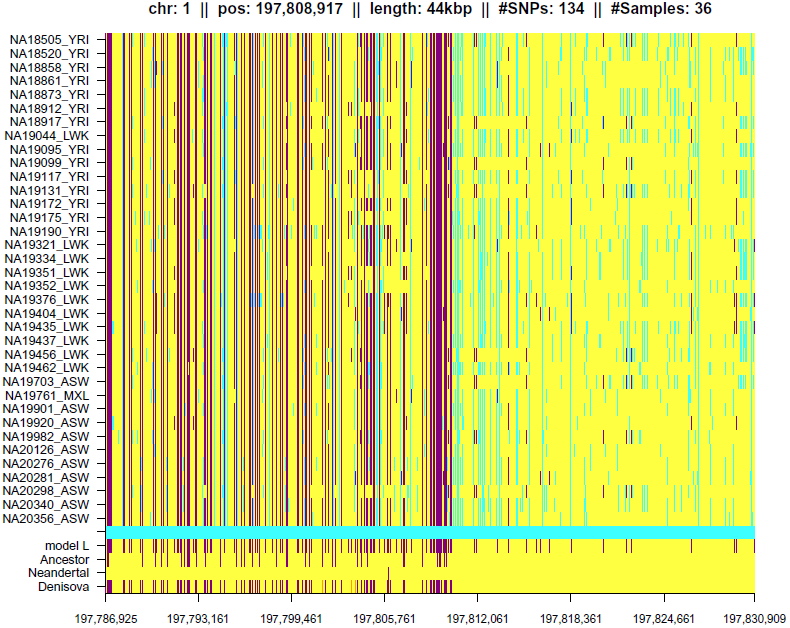
Example of an IBD segment matching the Denisova genome shared by Africans and one Admixed American. Only the first part of the IBD segment matches the Denisova genome and part of the IBD segment is shared by the majority of individuals. Many tagSNVs are also present in the reconstructed ancestor sequence, but they are not present in the Neandertal genome. See Figure 63 for a description.

**Figure 65:**
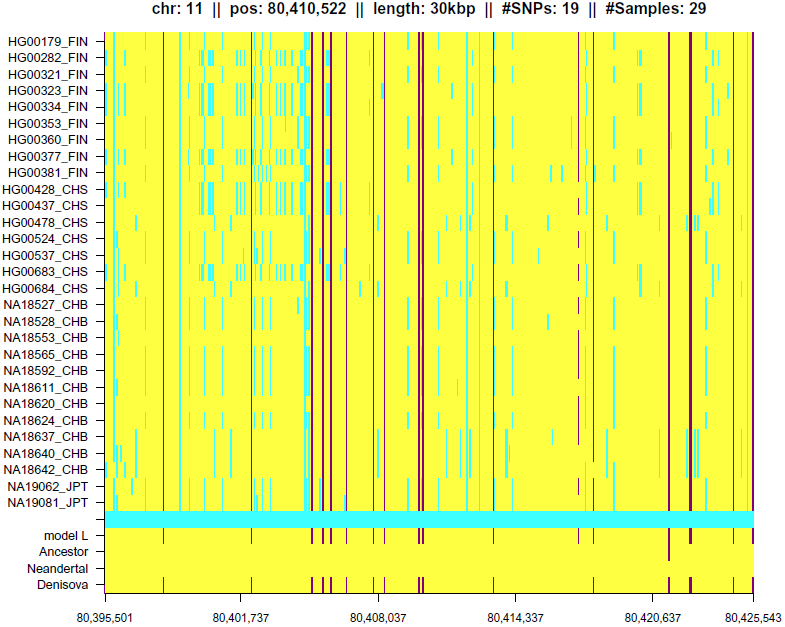
Example of an IBD segment matching the Denisova genome shared by Europeans and Asians. See Figure 63 for a description.

Figure 66 shows an example of an IBD segment that matches the Neandertal genome and is shared by Europeans, Asians, Admixed Americans, and Americans with African ancestry from SW US. Figure 67 shows an IBD segment that matches the Neandertal genome and is found exclusively in Asians.

**Figure 66:**
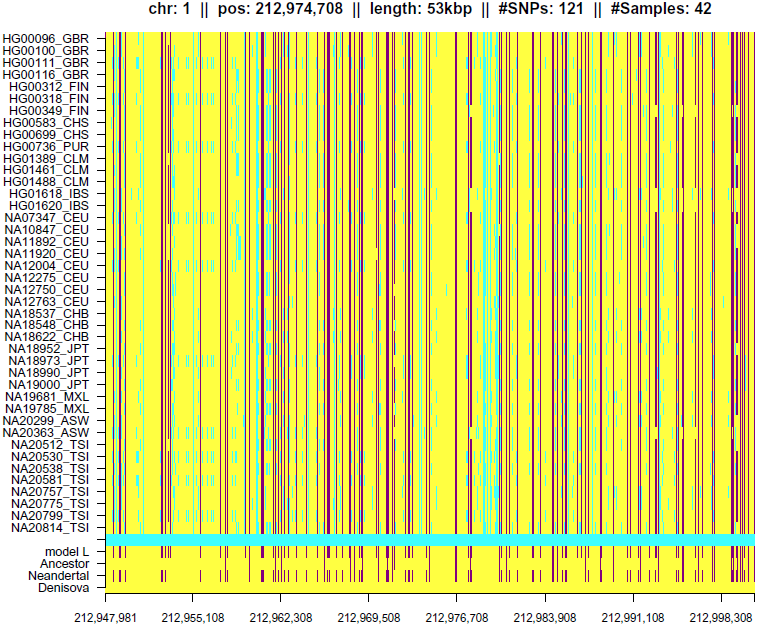
Example of an IBD segment matching the Neandertal genome shared by Europeans, Asians, Admixed Americans, and Americans with African ancestry from SW US. See Figure 63 for a description.

**Figure 67:**
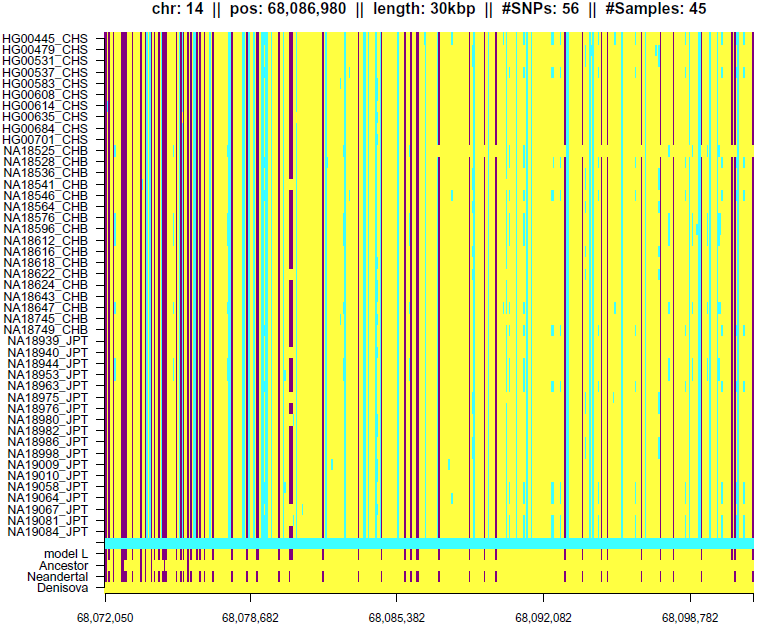
Example of an IBD segment matching the Neandertal genome shared exclusively among Asians. See Figure 63 for a description.

Figure 68 shows an example of an IBD segment matching the Neandertal and the Denisova genome shared by Asians, Admixed Americans, and Americans with African ancestry from SW US. Figure 69 shows an example of an IBD segment matching the Neandertal and the Denisova genome shared by Africans and Admixed Americans.

**Figure 68:**
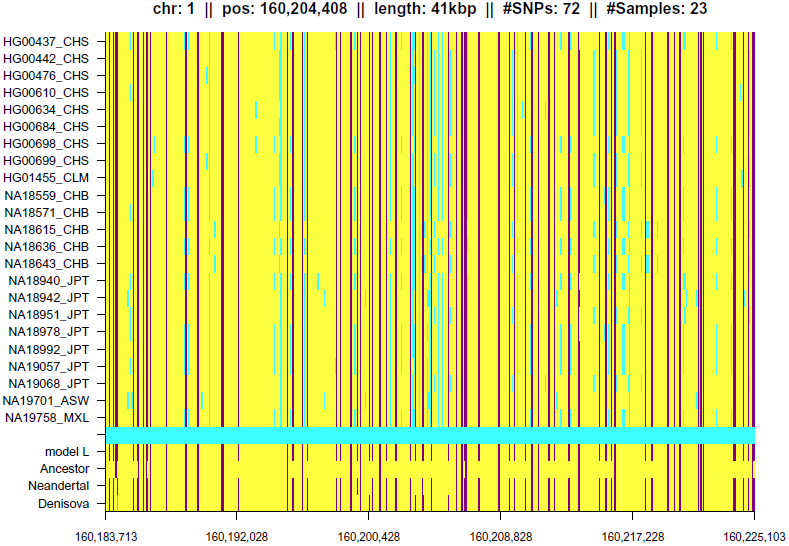
Example of an IBD segment matching the Neandertal and the Denisova genome shared by Asians, Admixed Americans, and Americans with African ancestry from SW US. See Figure 63 for a description.

**Figure 69:**
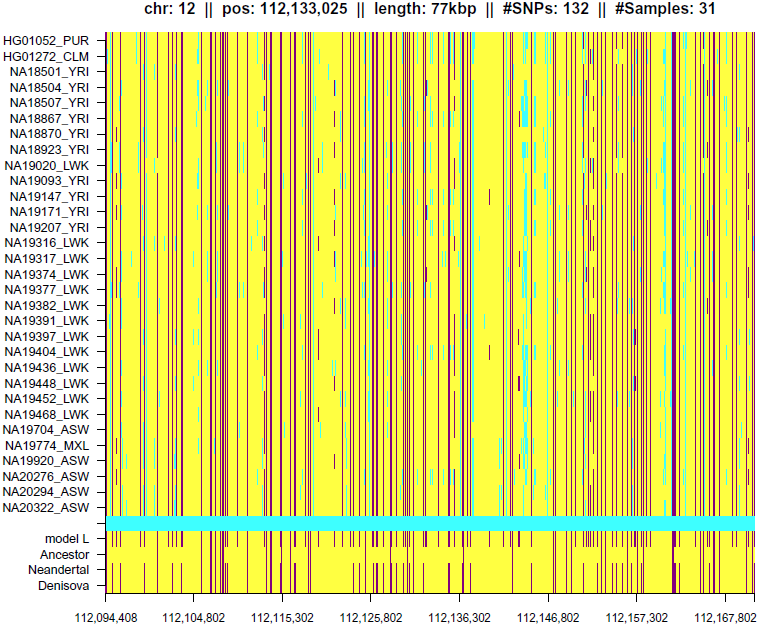
Example of an IBD segment matching the Neandertal and the Denisova genome shared by Africans and Admixed Americans. See Figure 63 for a description.

## 7 Conclusion

We have applied HapFABIA to the 1000 Genomes Project data to extract very short identity by descent (IBD) segments and analyzed the IBD sharing patterns of human populations, Neandertals, and Denisovans.

Some IBD segments are shared with (1) the reconstructed ancestral genome of humans and other primates, (2) the Neandertal genome, and/or (3) the Denisova genome. We could confirm an introgression from Denisovans into ancestors of Asians after their migration out of Africa. Furthermore, IBD sharing hints at a gene flow from Neandertals into ancestors of Asians and Europeans after they left Africa — to a larger extent into ancestors of Asians. Interestingly, many Neandertal- and/or Denisova-matching IBD segments are predominantly observed in Africans, some of them even exclusively. IBD segments shared between Africans and Neandertals or Denisovans are strikingly short, therefore we assume that they are very old. Consequently, we conclude that DNA regions from ancestors of humans, Neandertals, and Denisovans have survived in Africans. As expected, IBD segments on chromosome X are on average longer than IBD segments on the autosomes. Neandertal-matching IBD segments on the X chromosome confirm gene flow from Neandertals into ancestors of Asians and Europeans outside Africa that was already found on the autosomes. Interestingly, there is hardly any signal of Denisova introgression on the X chromosome.

In the near future the 1000 Genomes Project will be completed by genome sequences from additional populations, the UK10K project extends the subset of European genomes, and, most importantly, more ancient genomes will be successfully sequenced (Ust-Ishim, El Sidrón, etc.). Therefore, we expect that the analysis of the population structure by sharing patterns of very short IBD segments will become an increasingly important method in population genetics and its results will become more and more fine-grained.

## Appendices

### A Are Rare Variants Recent or Old?

Rare variants are found to be old which might seem to contradict results of other researchers.

However, the conclusion that rare variants are, in general, recent does not contradict our results. On the contrary, that the majority of IBD segments extracted by HapFABIA is found in Africans can explain why three times as many variants with 0.5–5% minor allele frequency are found in Africans as in Europeans or Asians in the 1000 Genomes Project (53). In particular, if rare variants are caused by a recent population growth, then the numbers found by the 1000 Genomes Project (53) are difficult to interpret. We want to discuss two reasons why rare variants may be old.

**(I)** Many rare variants are old, however, most rare variants may be recent, e.g. because of a recent growth in population. We did not consider private variants and have a bias toward less rare variants because, the more individuals share an IBD segment, the higher its significance, the more likely it is detected by HapFABIA. In our analysis, only 36% of the rare variants (not counting private variants) are in IBD segments. Therefore, the remaining majority of rare and private variants might be recent.

The vast majority of IBD segments is found in Africans. This would explain why Africans have more rare and low-frequency variants than Europeans or Asians. The publication of the 1000 Genomes Project (53) reports:

> “individuals from populations with substantial African ancestry (YRI, LWK and ASW) carry up to three times as many low-frequency variants (0.5–5% frequency) as those of European or East Asian origin,”

Table S14 in the supplementary information of the publication of the 1000 Genomes Project (53) provides the number of derived variants per individual in each population in its last rows (DAF denotes “derived allele frequency”, i.e. the frequency of the mutation):

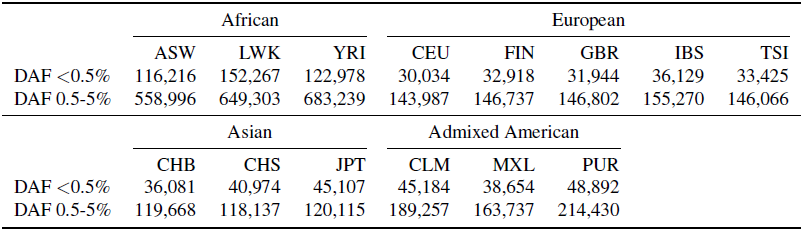

Recent variants are supposed to be derived variants. The table shows about four times more variants with derived allele frequency of 0.5-5% in Africans than in other populations (if we ignore the Admixed Americans that have African admixture). For derived allele frequency *<* 0.5%, we observe 2.5 times more variants in Africans than in Europeans or Asians.

**(II)** Many rare variants are old, however, compared to common SNPs, they are recent, i.e. the temporal relations remain, but variants are dated further back. We state in the manuscript that many rare variants are old and from times before humans migrated out of Africa. However, many common SNPs are even older and stem from common ancestors of human and chimpanzee. This means that many rare variants are old, but compared to common SNPs they are recent.

The fact that common SNPs are old is supported by findings of the Chimpanzee Consortium. In Hacia et al. (18) it was found that, of 397 human SNP sites, 214 were ancestral (shared with common ancestors of chimpanzee and human). Of the ancestral SNPs, 1/4 had the minor allele as ancestral allele. For the chimpanzee genome (54), it was found that

> “Of *∼*7.2 million SNPs mapped to the human genome in the current public database, we could assign the alleles as ancestral or derived in 80% of the cases according to which allele agrees with the chimpanzee genome sequence”

and that

> “a significant proportion of derived alleles have high frequencies: 9.1% of derived alleles have frequency *≥* 80%.”

According to (54, Suppl. Fig. S9) about 25% of the derived alleles have frequency *≥* 50% in which case the minor allele is the ancestral allele.

That some SNPs are very old and that some haplotypes are shared between humans and chimpanzee was also found in Leffler et al. (30):

> “We conducted a genome-wide scan for long-lived balancing selection by looking for combinations of SNPs shared between humans and chimpanzees. In addition to the major histocompatibility complex, we identified 125 regions in which the same haplotypes are segregating in the two species,”

In a recent publication (12), it was found that

> “The average age across all SNVs was 34,200 *±* 900 years (*±* s.d.) in European Americans and 47,600 *±* 1,500 years in African Americans, and these estimates were robust to sequencing errors […]”.

The authors further write

> “SNVs shared between European Americans and African Americans were significantly older (104,400 years and 115,800 years for European Americans and African Americans, respectively)”.

These figures are visualized as bar plots in Figure 70.

**Figure 70:**
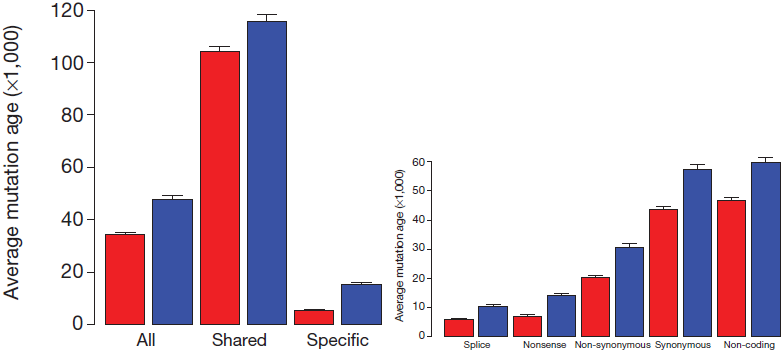
Left: Average age of all SNVs, “Shared” SNVs found both in European Americans (red) and in African Americans (blue), and SNVs found in only one population (“Specific”). Right: Average age for different functional types of variants. Both plots are taken from Fu et al. (12).

### B Population Groups of the 1000 Genomes Project

The subpopulations of the 1000 Genomes Project are given in Table 11 and the locations, where the individuals reside, are shown on the map in Figure 71.

**Table 11:**
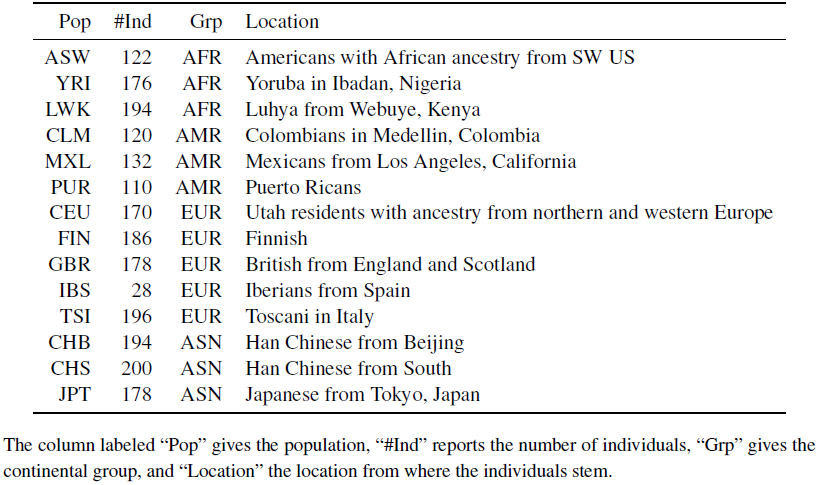
Overview of population groups of the 1000 Genomes Project.

**Figure 71:**
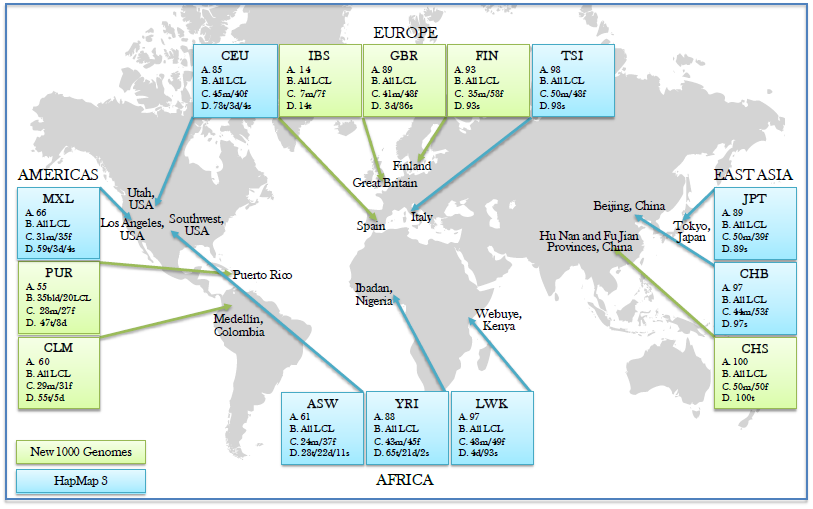
Map including the locations of the populations the 1000 Genomes Project phase I. Figure taken from (53).

## Acknowledgments

We thank Rokus van den Dool and others on several blogs for useful comments and discussions on an earlier draft of this manuscript.

